# SIPA1L1/SPAR1 is a non-PSD protein involved in GPCR signaling

**DOI:** 10.1101/2021.02.12.430872

**Authors:** Ken Matsuura, Shizuka Kobayashi, Kohtarou Konno, Miwako Yamasaki, Takahiro Horiuchi, Takao Senda, Tomoatsu Hayashi, Kiyotoshi Satoh, Fumiko Arima-Yoshida, Kei Iwasaki, Lumi Negishi, Naomi Yasui-Shimizu, Kazuyoshi Kohu, Shigenori Kawahara, Yutaka Kirino, Tsutomu Nakamura, Masahiko Watanabe, Tadashi Yamamoto, Toshiya Manabe, Tetsu Akiyama

## Abstract

SIPA1L1 (also known as SPAR1) has been proposed to regulate synaptic functions that are important in maintaining normal neuronal activities, such as regulating spine growth and synaptic scaling, as a component of the postsynaptic density (PSD)-95/*N*-methyl-D-aspartate receptor (NMDA-R)-complex. However, contrary to this view, our super-resolution and immunoelectron microscopic analyses demonstrate that SIPA1L1 is mainly localized to general submembranous and cytoplasmic regions in neurons, but scarcely to PSD. Our screening for native interactors of SIPA1L1 identified spinophilin and neurabin-1, regulators of G protein-coupled receptor (GPCR) signaling, but rejected PSD-95/NMDA-R-complex components. Furthermore, *Sipa1l1*^-/-^ mice showed normal spine size distribution and NMDA-R-dependent synaptic plasticity. Nevertheless, *Sipa1l1*^-/-^ mice showed aberrant responses to α_2_-adrenergic receptor (a spinophilin target) or adenosine A1 receptor (a neurabin-1 target) agonist stimulation and striking behavioral anomalies, such as hyperactivity, enhanced anxiety, learning impairments, social interaction deficits, and enhanced epileptic seizure susceptibility. Our findings revealed unexpected properties of SIPA1L1, suggesting a possible association of SIPA1L1 deficiency with neuropsychiatric disorders related to dysregulated GPCR signaling, such as epilepsy, attention deficit hyperactivity disorder (ADHD), autism, or fragile X syndrome.

## Introduction

Signal-induced proliferation-associated 1 (SIPA1)-like 1 (SIPA1L1) was identified as a PSD protein and a component of the PSD-95/NMDA-R complex in neurons (*1*). SIPA1L1 is comprised of a PSD-95/Dlg/ZO-1 (PDZ) domain, actin-interacting domains, a coiled-coil domain, and a GTPase-activating protein (GAP) domain specific to the Rap family of small GTPases. SIPA1L1 was shown to possess actin-reorganizing activity through its actin-interacting domains and that it promotes dendritic spine growth in cultured neurons (thus the name Spine-associated RapGAP, SPAR) (*1*). It was further reported that SIPA1L1 is bound and phosphorylated by Polo-like kinase 2 (Plk2) and targeted for degradation by the ubiquitin-proteasome pathway (*2*). Plk2-dependent degradation of SIPA1L1 is suggested to underlie the molecular mechanism of synaptic scaling (*3*). Degradation of SIPA1L1 and the resulting weakening of synapses is postulated to accompany shrinkage of dendritic spines and reduction of the number of surface AMPA-Rs (α-amino-3-hydroxy-5-methyl-4-isoxazolepropionic acid receptors) and to operate as a part of the small GTPase Ras and Rap signaling regulatory system in homeostatic synaptic plasticity (*4*).

SIPA1L1 was also shown to bind other proteins, including EphA4 receptor and the leucine zipper tumor suppressor (LZTS) family of proteins. EphA4 binds the PDZ domain of SIPA1L1 and is involved in neuronal cell adhesion or axonal growth cone morphogenesis through regulation of Rap1 activity (*5*). The LZTS family of proteins bind SIPA1L1 via a reciprocal coiled-coil domain interaction (*6, 7*). LZTS1/PSD-Zip70 has been suggested as critical in the spine localization of SIPA1L1, collaborating with SIPA1L1 for spine maturity and maintenance (*8*).

Spinophilin (also known as neurabin-2/PP1R9B) and its paralog, neurabin-1/PP1R9A, are F-actin-binding proteins enriched in dendritic spines (*9-11*). Spinophilin and neurabin-1 share similar domain structures, which comprise an F-actin-binding domain, a protein phosphatase 1-binding domain, a PDZ domain, and coiled-coil domains. A notable feature of this family of proteins is their ability to modulate GPCR signaling, which controls various physiological responses. The major difference between the two proteins is their binding and regulatory specificity for GPCRs via their low conserved receptor-binding domains. To date, spinophilin has been shown to target α_1_- and α_2_-adrenergic receptors (αARs) (*12*), muscarinic-acetylcholine receptors (mAchRs) (*13*), dopamine D2 receptors (*14*), μ-opioid receptors (*15*), and group 1 metabotropic glutamate receptors (Gp1 mGluRs) (*16*), whereas neurabin-1 targets adenosine A1 receptors (*17*).

Despite various important roles suggested for SIPA1L1 in neurons, its physiological role remains poorly understood. Here, we examined localization of SIPA1L1 in mature brain using super-resolution microscopy (SRM) and immunoelectron microscopy (IEM). Unexpectedly, we found that SIPA1L1 is generally localized submembranously in somata and neurites of neurons, and to cytoplasm in dendritic spines, but that it is scarce in PSD regions. Screening for native SIPA1L1 interactors in the mouse cerebrum validated spinophilin and neurabin-1 along with other candidate proteins. Finally, we addressed physiological functions of SIPA1L1 by histological, electrophysiological, pharmacological, and behavioral analyses of *Sipa1l1*^-/-^ mice. The results demonstrated a critical role of SIPA1L1 in certain types of GPCR signaling and in brain functions that are highly relevant to neuropsychiatric disorders.

## Results

### SIPA1L1 localizes to submembranous regions in neurons, but is scarcely associated with PSD

To investigate physiological roles of SIPA1L1, we generated mice lacking *Sipa1l1* (Fig.S1). We first examined the expression pattern and localization of SIPA1L1 in mature brain. *Sipa1l1* promoter activity was present throughout the brain, with the highest activity in the cerebrum, including the hippocampus, cerebral cortex, striatum, and olfactory bulb (Fig.1A). β-Gal staining was observed in most of NeuN positive neurons as well as NeuN negative neurons such as Purkinje cells in cerebellum, but not in glial cells (Fig. S2A-D). The results suggested *Sipa1l1* expression is mostly neuron-specific in the brain, if not neuron-exclusive. The strong immunofluorescence signal of SIPA1L1 was detected in wild-type (WT) cerebrum (Fig. 1B), consistent with the pattern of *Sipa1l1* promoter activity, but not in the *Sipa1l1*^-/-^ brain (Fig. 1C). In the hippocampus, SIPA1L1 immunoreactivity had a relatively stronger signal in the CA1 region, with strong signals in both somata and neuropil regions (Fig. 1D, E). SIPA1L1 was not only expressed in excitatory neurons, but also expressed in virtually all GABAergic neurons observed (Fig. S2E-G).

**Figure 1.**
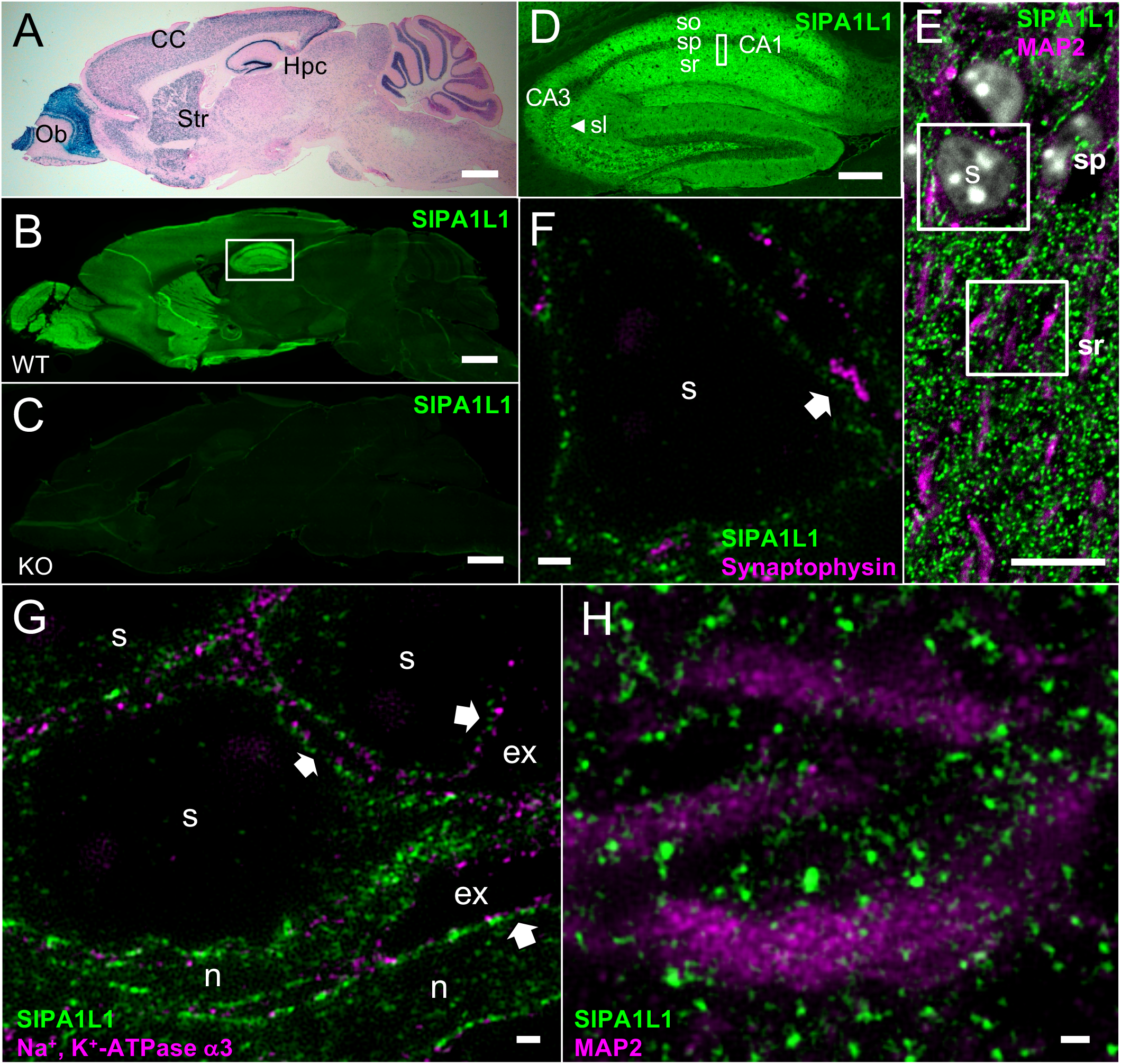
SIPA1L1 localizes to submembranous regions in the soma of neurons in mature brain. (**A**) X-Gal staining of *Sipa1l1*^+/-^ brain. (**B**-**D**) Immunostaining of wild-type (B and D) or *Sipa1l*^-/-^ (C) brain using anti-SIPA1L1 antibody. (D) is an image corresponding to the boxed area in (B). (**E**-**H**) Double immunostaining images of SIPA1L1 and MAP2 (E and H), synaptophysin (F) or Na^+^, K^+^-ATPase α3 (G). (E) is an image corresponding to the boxed area in (D). (F and G) or (H) are images corresponding to the top or bottom boxed area in (E), respectively. Note that higher magnification images were obtained independently and are not a direct magnification of boxed areas. (B-D), (E), or (F-H) are fluorescent, confocal, or super-resolution microscopic images, respectively. Arrows in (F) and (G) show examples of submembranous distributions of SIPA1L1. The arrow in (F) also indicates a presynaptic terminal represented by synaptophysin. CC, cerebral cortex; Hpc, hippocampus; Ob, olfactory bulb; Str, striatum; so, stratum oriens; sp, stratum pyramidale; sr, stratum radiatum; sl, statum lucidum, s, soma; ex, extracellular area; n, proximal neurite. Scale bars:: (A-C) 1000 μm; (D) 200 μm; (E) 10 μm; (F-H) 1 μm.

We next minutely investigated the subcellular localization of SIPA1L1, utilizing a confocal-based spinning disk super-resolution microscope, which implements structured illumination microscopy (SIM) and achieves a spatial resolution of 120 nm with high scanning speed (*18*). SIPA1L1 was primarily distributed beneath the plasma membrane in somata and in proximal neurites of pyramidal neurons in the CA1 region (Fig. 1F, G). SIPA1L1 was relatively evenly distributed, and no specialized structure or distribution was observed in regions apposing presynaptic terminals (Fig. 1F). In the neuropil region, co-staining of SIPA1L1 with dendritic marker MAP2 showed large clusters of SIPA1L1 surrounding the dendritic shaft and also smaller signals embedded within the shaft (Fig. 1H). Quadruple staining of SIPA1L1, bassoon, synapsin-1/2, and F-actin in the neuropil region suggested the generally postsynaptic localization of SIPA1L1 (Fig. 2A), consistent with the previous report in cultured neurons (*1*).

**Figure 2.**
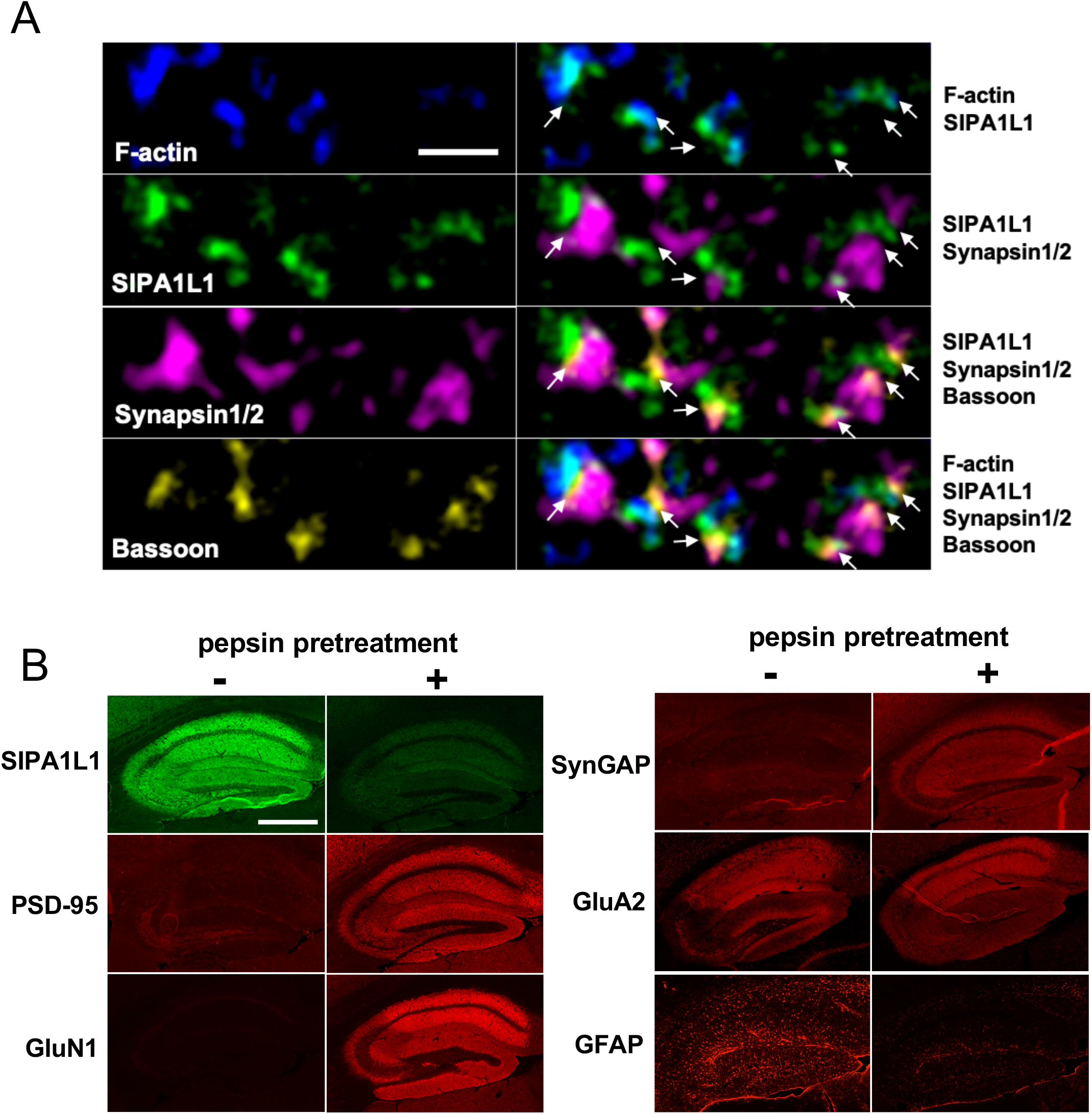
SIPA1L1 localizes to postsynapses but shows non-PSD-like staining pattern. **(A)** Super-resolution immunofluorescence images of SIPA1L1, F-actin, synapsin1/2, and bassoon in the hippocampal CA3 stratum lucidum. Synapsin1/2 or bassoon represents presynaptic vesicles or presynaptic active zone, respectively. F-actin is a major cytoskeletal structure in the postsynaptic spine head. Arrows point to presynaptic active zones (bassoon) between the opposing SIPA1L1 and synapsin1/2 staining. (**B**) Immunofluorescent images of indicated proteins on pepsin-pretreated (+) or -untreated (-) hippocampal slices. Paired images are acquired with identical settings and conditions. PSD proteins show strong staining with pepsin pretreatment. GluA2 is known to have both PSD and non-PSD populations (Ref. 29). Scale bars: (A) 1 μm; (B) 500 μm.

We further proceeded to confirm that SIPA1L1 associates with PSD. It has been shown that conventional immunostaining methods fail to show the true distribution of PSD proteins in brain tissue due to the densely packed nature of PSD, so unmasking of epitopes such as by protease pretreatment is required (*19*). Accordingly, representative PSD proteins, such as PSD-95, NMDA-R subunits, and SynGAP, a downstream signaling protein that binds to PSD-95, all showed strong specific staining only after pepsin pretreatment. However, SIPA1L1 and the non-PSD protein, glial fibrillary acidic protein (GFAP), showed strong staining without antigen unmasking, and their signals decreased significantly after pepsin pretreatment (Fig. 2B). These results suggested that a significant proportion of SIPA1L1 is not in a densely packed PSD and it localizes to regions to which antibodies and pepsin have easy access, facilitating the detection by an antibody and the degradation by pepsin.

To investigate in more detail, we performed IEM in hippocampal CA1 neuropil region, using *Sipa1l1*^-/-^ brain tissue as a negative control. The PSD in electron micrograph is defined as electron-dense structure extending 30-50 nm into the cytoplasm beneath the postsynaptic membrane. Accordingly, PSD-95 staining showed distribution mostly within 40 nm from the midline of synaptic cleft. Surprisingly, however, SIPA1L1 staining showed a broad distribution within 60 - 200 nm from the midline of synaptic cleft, peaking around 120 nm, but scarce staining within 0 - 60 nm (Fig. 3A and B). This showed striking contrast to DAP/GKAP, a protein that binds directly to PSD-95 at the same domain that binds SIPA1L1, which shows clear peak within PSD area by IEM (*20*). This result indicated that the vast majority of SIPA1L1 is not in close proximity to PSD-95, as would occur in direct binding. Predominant postsynaptic localization of SIPA1L1 was also confirmed by IEM. (Fig. 3C)

**Figure 3.**
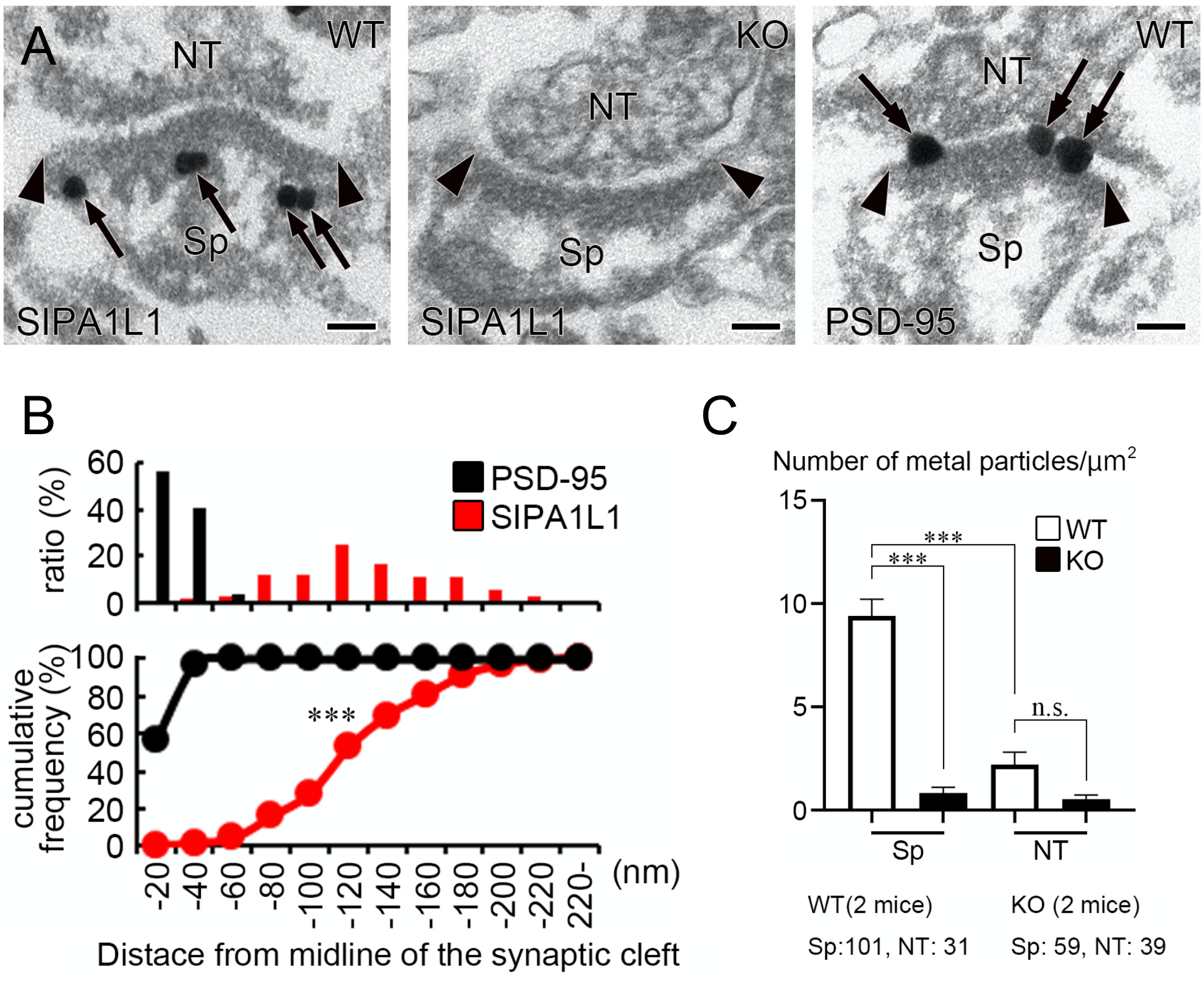
SIPA1L1 localizes to non-PSD region in dendritic spines. **(A)** Electron micrographs of hippocampal CA1 stratum radiatum labeled for SIPA1L1 and PSD-95. *Sipal11*^-/-^ (KO) brain was used for negative control. The arrows show immuno-metal particles, and the arrowheads indicate the extent of PSD. Sp, dendritic spine; NT, presynaptic nerve terminal. Scale bar: 100 nm. (**B**) Distribution of SIPA1L1 and PSD-95 in dendritic spines shown as distance from midline of the synaptic cleft. Data are put into 20 nm bins. Total of 148 and 196 metal particles for SIPA1L1 and PSD-95, respectively, were analyzed. Two tailed Kolmogorov-Smirnov test, ***P < 0.0001. (**C**) Density of immuno-metal particles labeled for SIPA1L1 was measured for dendritic spines and presynaptic nerve terminals, respectively, in WT and SIPA1L1 KO hippocampus. Total number of metal particles analyzed are shown below the graph. Mean ± SEM is shown. One-way ANOVA followed by Tukey’s post-hoc test, ***P < 0.0001.

These non-PSD or submembranous localizations of SIPA1L1 suggest an extrasynaptic and/or more general function of SIPA1L1 in neurons, which has not been appreciated.

### *Sipa1l1*^*-/-*^ mice show normal spine size distribution and NMDA-R-dependent synaptic plasticity

As exogenous SIPA1L1 expression was shown to promote spine head growth (*1*), and its Plk2-dependent degradation is suggested to result in spine shrinkage in hippocampal neuronal cultures (*3, 4*), we examined the change in cross-sectional areas of spine heads and PSD lengths in the CA1 stratum radiatum of *Sipa1l1*^-/-^ hippocampus, using electron microscopy. These two parameters represent the volume of spines; hence, they can be used to deduce changes in spine size (*21*). Gross ultrastructural features of asymmetric glutamatergic synapses, synaptic density, and global distribution of spine head area or PSD length in *Sipa1l1*^-/-^ mice, were all comparable to those of WT mice (Fig. 4A-D). These results suggested that SIPA1L1 is generally dispensable in spine growth and maturation.

**Figure 4.**
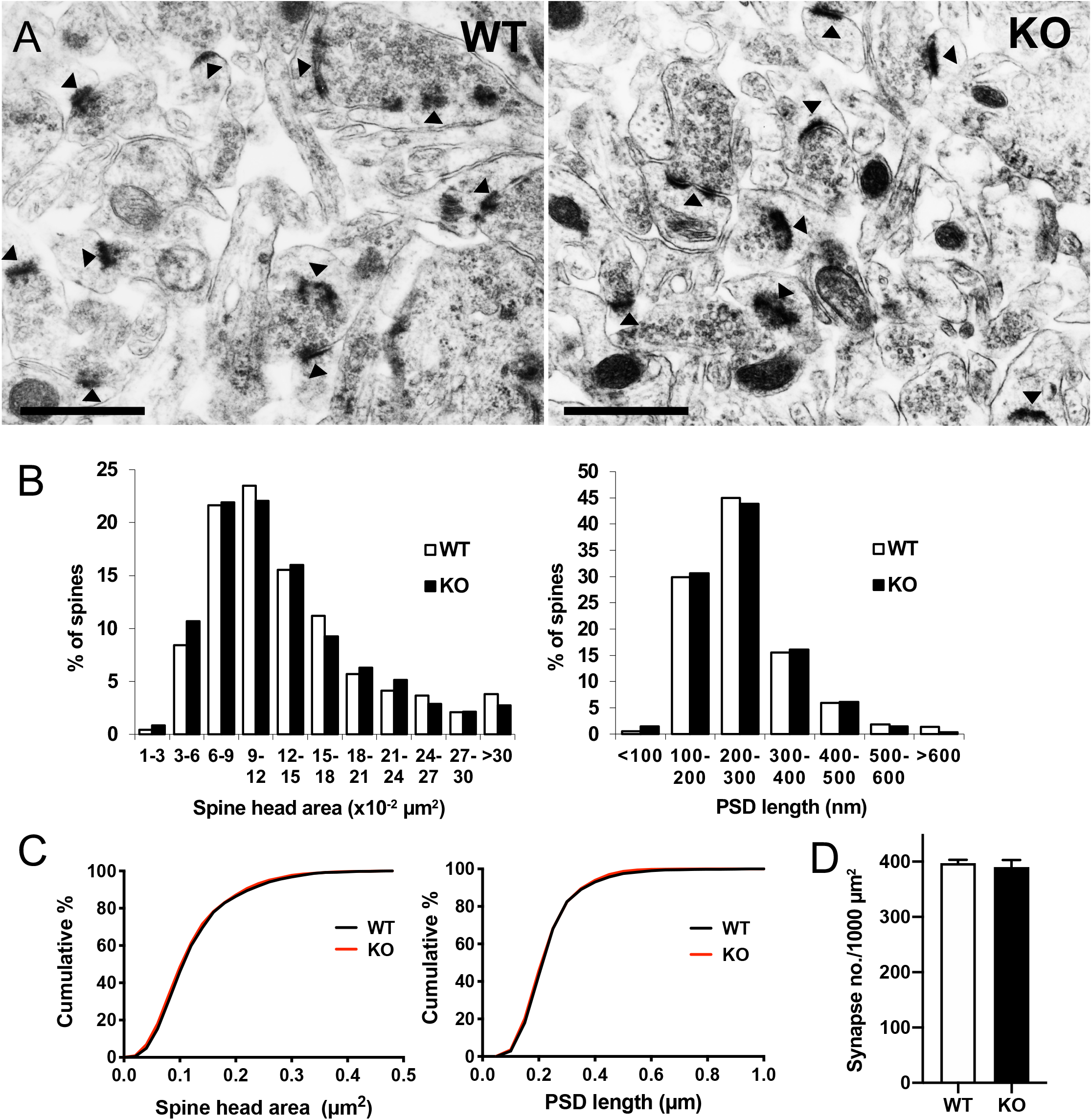
*Sipa1l1*^*-/-*^ mice show normal synaptic density and spine size distributions. (**A**) Representative thin-section electron micrographs of hippocampal CA1 stratum radiatum, showing normal asymmetric morphology of synapses in both WT and KO mice. Arrowheads point to PSD in dendritic spines. Scale bar: 1 μm. (**B**) Frequency or (**C**) cumulative distribution plots of cross-sectional spine head area and PSD length. N = 2110 (WT) or 2107 (KO) spines from 4 mice per genotype. Two tailed Kolmogorov-Smirnov test, D = 0.034, p = 0.17 or D = 0.030, p = 0.29 for spine head area or PSD length, respectively. (**D**) Synaptic density calculated from same electron micrographs as (B) and (C). Total 2110 (WT) or 2107 (KO) synapses were analyzed. Mean ± SEM is shown. t (6) = 0.512, P = 0.63; unpaired two-tailed t test.

We also performed electrophysiological experiments to address SIPA1L1 deficiency in synaptic transmission and NMDA-R-dependent plasticity. Paired-pulse facilitation (PPF) and input–output relationships of excitatory postsynaptic potentials (EPSPs) in the hippocampal CA1 stratum radiatum showed no differences between *Sipa1l1*^-/-^ and WT mice (Fig. 5A and B). This is consistent with the postsynaptic localization of SIPA1L1 and the similar spine size/density that exists between the genotypes. This result also suggests that no general depression of the AMPA-R mediated response occurred in *Sipa1l1*^-/-^ hippocampus, which could have been resulted if Rap signaling was constitutively activated (*22, 23*). NMDA-R-dependent LTP induced by high-frequency stimulation was also comparable between the genotypes (Fig. 5C). Thus, SIPA1L1 may also be dispensable in actin reorganization and dynamic changes of spine morphology that underlie synaptic plasticity in hippocampal CA1 LTP (*24*).

**Figure 5.**
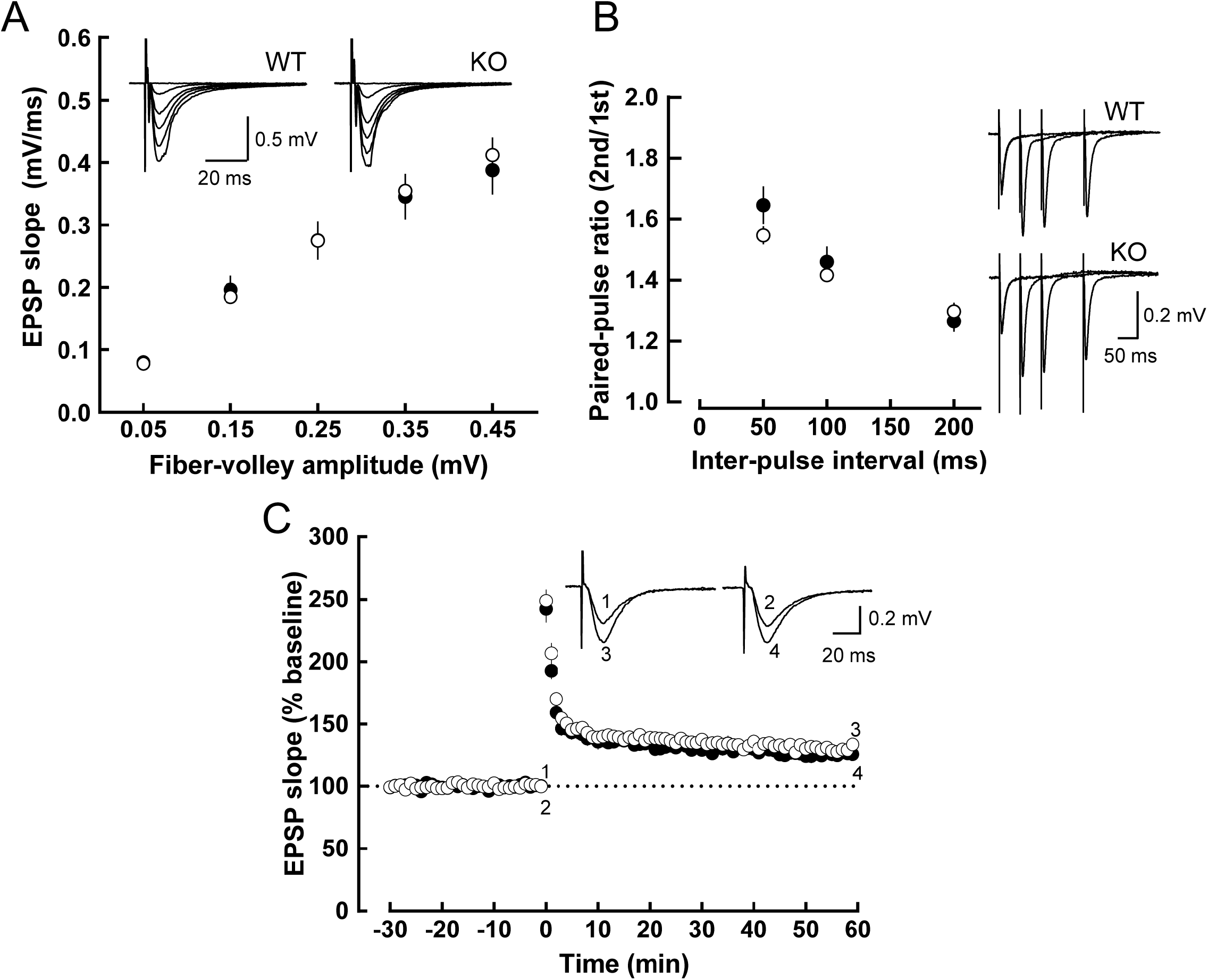
Basal synaptic transmission, paired-pulse facilitation, and hippocampal CA1 LTP are normal in *Sipa1l1*^*-/-*^ mice. **(A)** The input (fiber-volley amplitude)-output (EPSP slope) relationship of AMPA receptor-mediated EPSPs at Schaffer collateral-CA1 pyramidal cell synapses in acute hippocampal slices of WT (open circles: n = 10) and *Sipa1l1*^-/-^ (KO, closed circles: n = 9) mice. There was no significant difference between two genotypes. Sample traces of EPSPs (average of 10 consecutive sweeps) evoked with various stimulus strengths are shown in the inset. (**B**) Paired-pulse facilitation. The paired-pulse ratio (the ratio of slopes of second EPSPs to those of first EPSPs) is shown as a function of inter-pulse intervals (IPI) in the presence of 25 mM D-AP5. In any IPI (50, 100, and 200 ms), no significant difference was observed between WT (open circles, n = 6 slices) and *Sipa1l1*^-/-^ (closed circles, n = 6 slices) mice. Right panel: sample traces of synaptic responses evoked by paired stimuli at intervals of 50, 100 and 200 ms are superimposed. (**C**) The time course of LTP induced by tetanic stimulation in WT (open circles, n = 13 slices) and *Sipa1l1*^-/-^ (closed circles, n = 12 slices) mice. A train of high-frequency stimuli (100 Hz, 1 s) was delivered at time 0. Sample traces (average of 10 consecutive responses) in the inset were EPSPs obtained at times indicated by the numbers in the graph.

We next considered the possibility of compensation by up-regulation of SIPA1L1 homologs. SIPA1L1 has two paralogs, denominated SIPA1L2 and SIPA1L3. Both proteins are localized to the postsynaptic compartment and interact with the LZTS family of proteins, similar to SIPA1L1 (*25, 26*). However, SIPA1L2 does not colocalize with F-actin when expressed in COS-7 cells nor induce spine growth in primary cultured neurons (*25*). We found that the expression of SIPA1L2 and SIPA1L3 was relatively higher in the hippocampus compared to other regions in WT brain, which was especially prominent for SIPA1L3 (Fig. S3A-D). However, we did not observe any significant change in expression level of SIPA1L2 or SIPA1L3 in *Sipa1l1*^-/-^ hippocampus compared to WT (Fig. S3E and F). These results suggested little contribution of SIPA1L1 paralogs to compensation for SIPA1L1 deficiency.

### Screening for native SIPA1L1 interactors in the brain identified spinophilin, neurabin-1, and drebrin

To find a clue for physiological function of SIPA1L1 and to clarify the discrepancy between the observed non-PSD localization of SIPA1L1 and reported SIPA1L1-interacting PSD-associated proteins, we performed screening for physiological SIPA1L1-interacting proteins. We adopted a chemical crosslinking immunoprecipitation (cIP) strategy to preserve native interactions prior to addition of detergents. This strategy also enabled us to use stringent solubilization (2% SDS) and wash conditions to minimize nonspecific or artifactual interactions.

We performed cIP combined with mass spectrometry (cIP-MS) screening in the mouse cerebral cortex and hippocampus, and identified 120 candidate SIPA1L1-interacting proteins (Fig. 6A and Table S1). We were able to successfully validate these interactions using cIP-Western blotting (cIP-WB) on high-ranking proteins, which were mostly chosen based on mutual detection in both brain regions as general interactors (Fig. 6B). These included known SIPA1L1-binding proteins, such as α-actinin-1, LZTS1/PSD-Zip70, LZTS3/Pro-SAPiP1, and notably, novel interactors spinophilin/PP1R9B, neurabin-1/PP1R9A, and drebrin. On the other hand, reported SIPA1L1 interactors including those that are strongly associated with PSD, namely, PSD-95, SynGAP, and neuroligin-1 failed to be reliably detected in cIP-WB (Fig. 6B). A parallel experiment on cIP-WB using an anti-PSD-95 antibody for IP resulted in successful detection of well-established PSD-95 interactors, such as GluN1, GluN2B, SynGAP, and DAP/GKAP, but not SIPA1L1 (Fig.6C). These results are consistent with the scarce localization of SIPA1L1 to PSD as indicated by IEM. The pepsin pretreatment-immunostaining analysis showed that the SIPA1L1-interacting proteins have stronger staining in pepsin-untreated brain slice, similar to SIPA1L1 (Fig. S4). This is in line with reports showing their cytoplasmic localization in the dendritic spine, although some of these proteins also localize to PSD (*27-29*).

**Figure 6.**
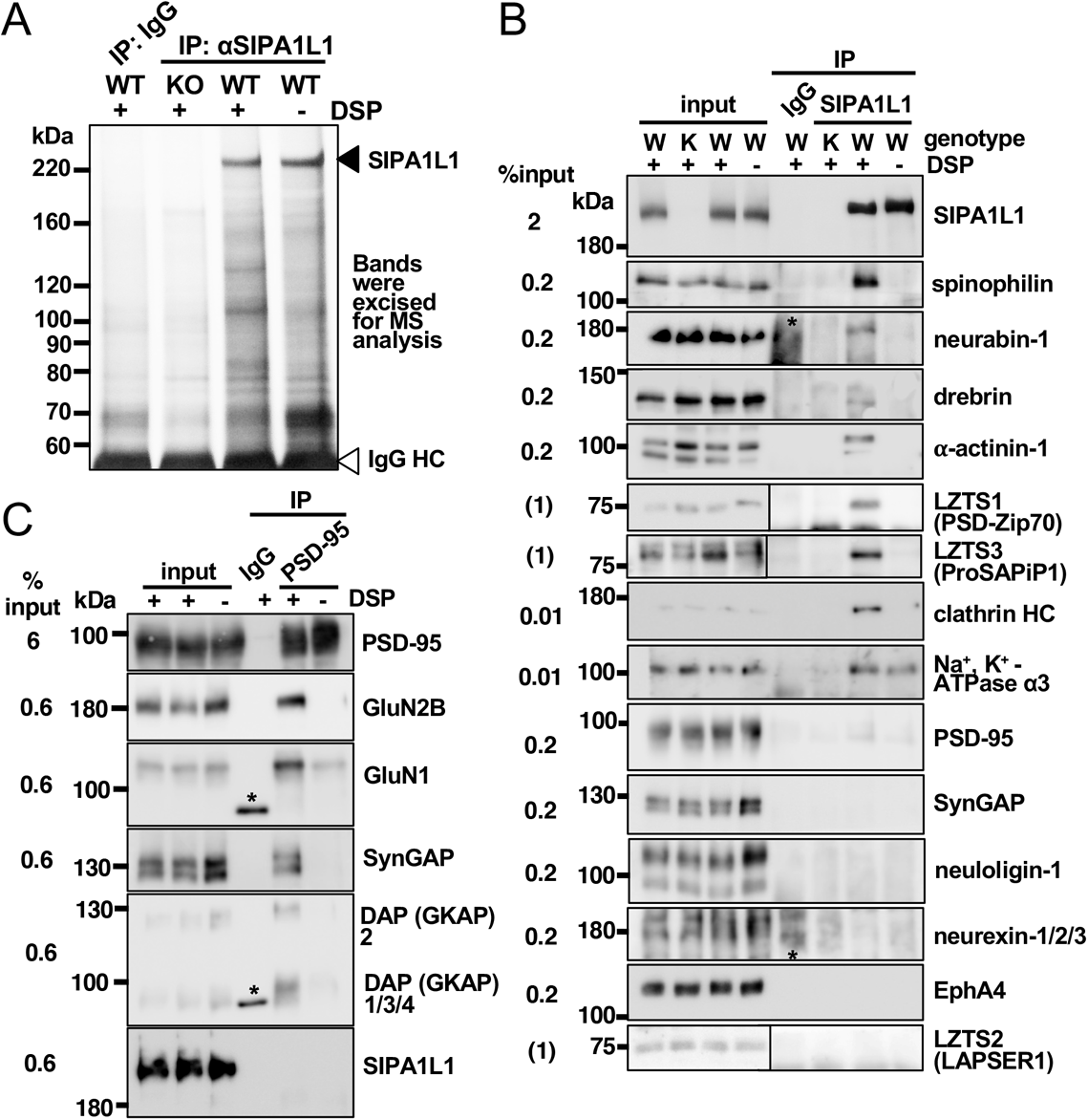
Identification of native interactors of SIPA1L1 in the brain. **(A)** Silver staining analysis of an SDS-PAGE gel showing the recovery of proteins co-precipitated with SIPA1L1 or IgG controls from cerebral cortex lysate. Note bands specific for SIPA1L1 expression and DSP crosslinking. See Table S1 for bands excised and results of LC-MS/MS analyses of these bands. (**B**) Confirmation of MS results by cIP-WB. The proportion of clathrin HC or Na^+^, K^+^-ATPase α3 co-precipitated with SIPA1L1 relative to the whole population was much smaller compared to other interactors. Detection of inputs for the LZTS family of proteins was performed separately with an increased amount, as they were not detectable with 0.2% input. The detection of a protein band in a control non-crosslinked lane, such as is seen in Na+, K+-ATPase α3, may suggest artifactitious post-solubilization interaction. (**C**) The same experiment was performed as in (B) except that anti-PSD-95 antibody was used for immunoprecipitation. *indicates non-specific band.

We attained additional confirmation by super-resolution colocalization analysis using Confined Displacement Algorithm (Fig. S5) (*30*). Analyses in the neuropil of mouse hippocampus or cerebral cortex revealed highest colocalization of SIPA1L1 with spinophilin among the examined candidates (Fig. 7 and Table S2). Relatively high correlation of spinophilin and SIPA1L1 signals suggested constant stoichiometric ratio in a complex. Similar results were observed for two different antibodies raised against different regions of spinophilin (Table S2). Analysis of somata and proximal neurites of CA1 pyramidal neurons showed that spinophilin aligns with SIPA1L1 along submembranous regions with occasional co-localization (Fig. 8; compare with Fig. 1G). Neurabin-1 and drebrin also showed significant colocalization and correlation with SIPA1L1 in the cerebral cortex (Fig. 7). However, colocalization of SIPA1L1 with neurabin-1 in hippocampus was much lower (Manders coefficient M2 = 0.062) compared to cerebral cortex (M2 = 0.179) and not significantly correlated (Table S2). This may explain the reason why neurabin-1 was detected in cerebral cortex by cIP-MS but not in hippocampus. α-actinin-1 also showed significant colocalization with SIPA1L1 albeit overlaps were quite small and without correlation. Clathrin heavy chain, Na^+^, K^+^-ATPase α3, or synaptophysin, a presynaptic marker, did not show significant colocalization or correlation with SIPA1L1 (Fig. 7, Fig. S5, and Table S2).

**Figure 7.**
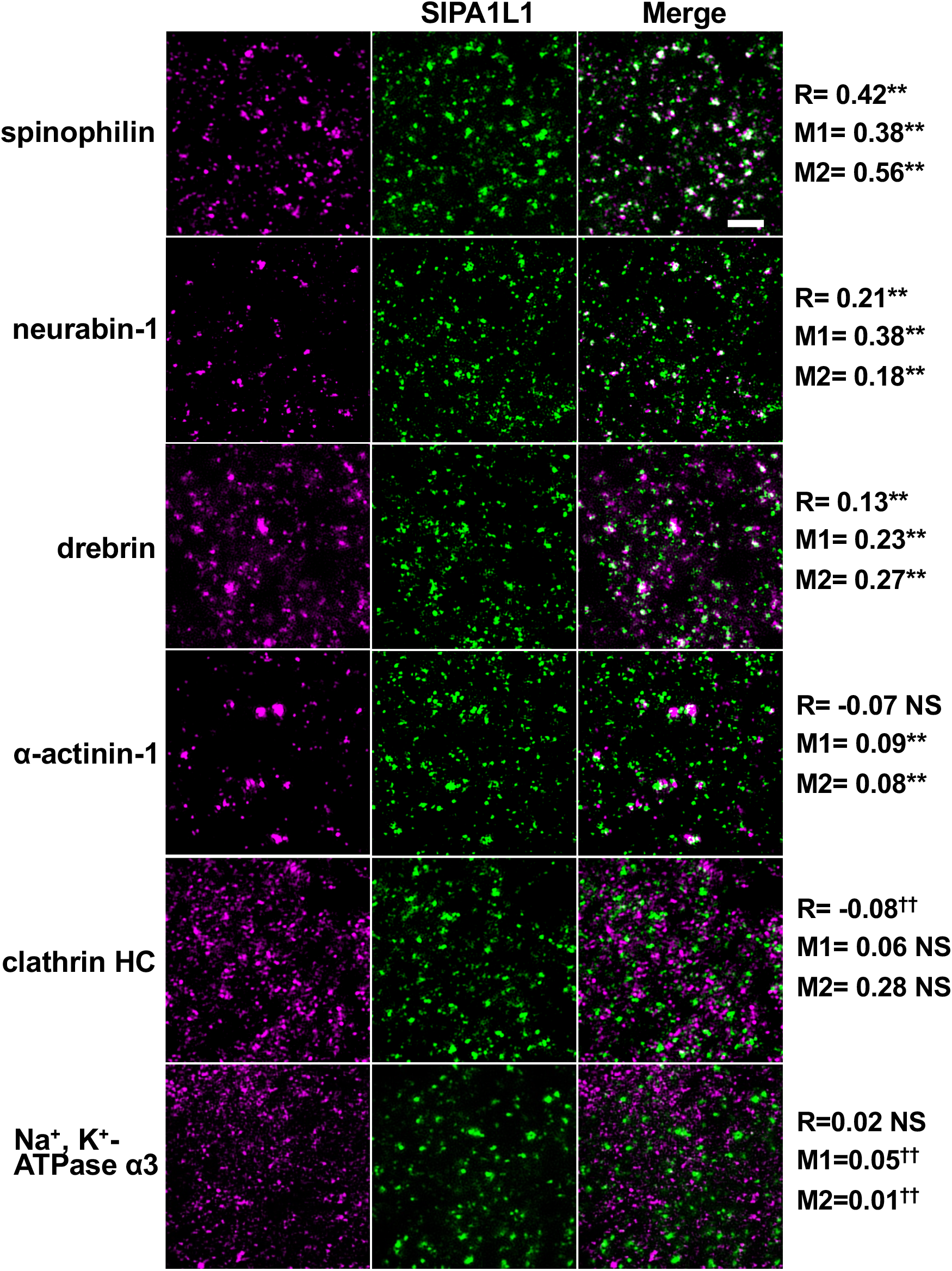
Spinophilin highly colocalizes with SIPA1L1 in the neuropil area of cerebrum. Representative super-resolution immunofluorescence images of SIPA1L1 co-stained with indicated proteins in the neuropil of layer V of the cerebral cortex. Colocalization was analyzed using Confined Displacement Algorithm on total 6724 μm^2^ of neuropil area per protein pair (see Fig. S5, Table S2, and Methods for details). R, Pearson correlation coefficient; M1 and M2, Manders coefficient for candidate interacting protein and SIPA1L1, respectively. **P < 0.01 (significant correlation or colocalization); ^**††**^ P < 0.01 (significant noncorrelation or noncolocalization compared to random displacement images); NS, not significant. Scale bar: 2 μm.

**Figure 8.**
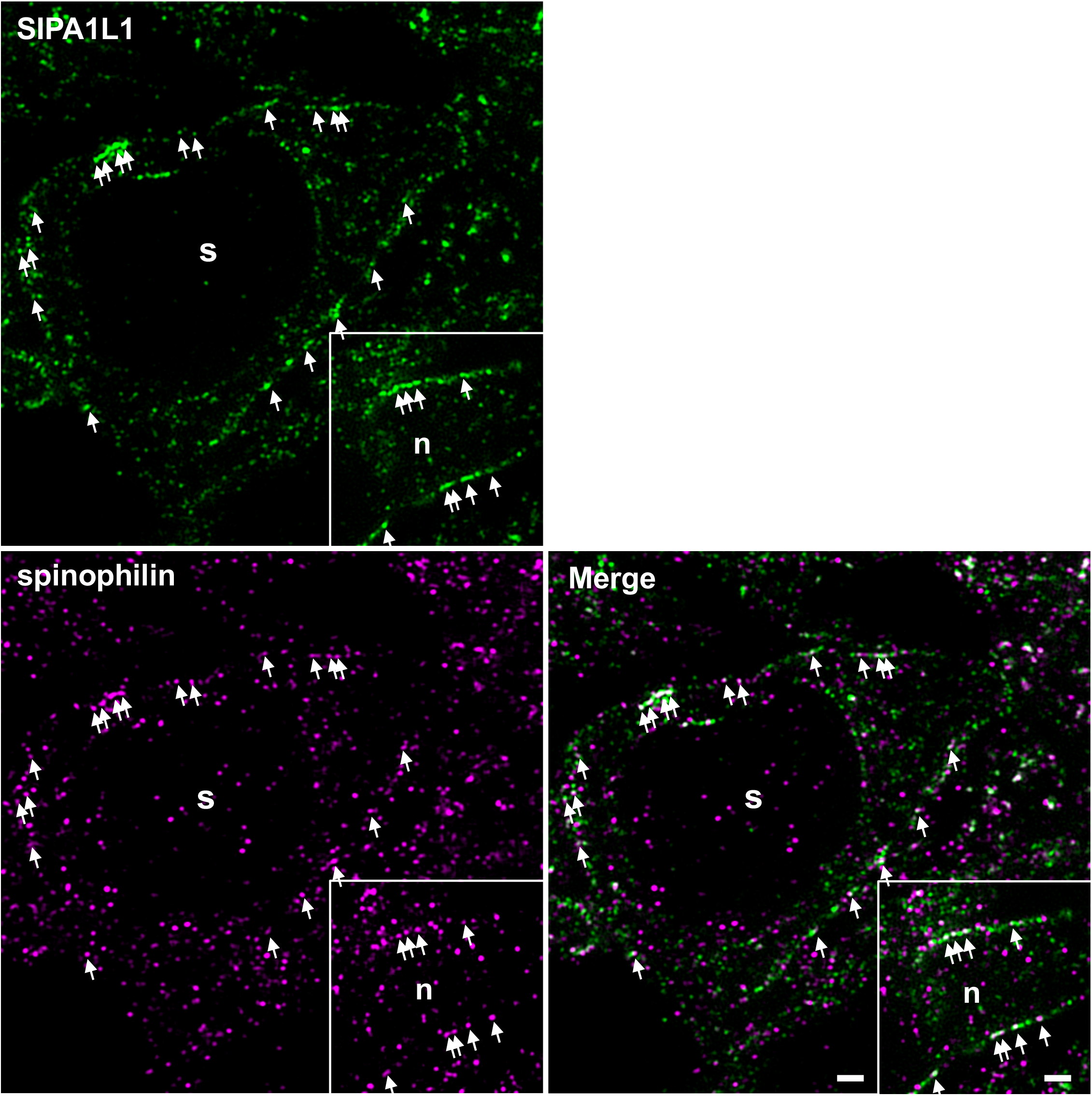
Spinophilin co-localizes with SIPA1L1 at the submembranous region in the soma of neurons. Representative super-resolution immunofluorescence images of SIPA1L1 co-stained with spinophilin in a soma and a proximal neurite (inset) of a pyramidal neuron in the hippocampal CA1 area. Arrows indicate examples of spinophilin aligned and colocalized with SIPA1L1 in the submembranous region (compare with Fig.1G). s, soma; n, proximal neurite. Scale bar: 1 μm.

We further confirmed that exogenous expression of SIPA1L1 and spinophilin in COS-7 cells successfully reproduced the SIPA1L1-spinophilin interaction without using a crosslinker. The interaction depended on the C-terminal coiled-coil domain, but not on the N-terminal F-actin-binding domain of spinophilin. This result suggested that the SIPA1L1-spinophilin interaction is not mediated by F-actin (Fig. S6).

Taken together, our screening identified spinophilin as the most promising physiological interactor of SIPA1L1 in the brain, and also neurabin-1 and drebrin as strong candidates. LZTS1 and LZTS3 may also be *bona fide* interactors as they showed clear and high proportion of co-IP with SIPA1L1 (Table S3 summarizes the results of the entire screening).

### *Sipa1l1*^*-/-*^ mice show aberrant responses to GPCR agonist stimulation and significantly enhanced epileptic seizure susceptibility

We next sought the functional relevance of the spinophilin-SIPA1L1 interaction. One of the most well-studied GPCR targets for spinophilin is α_2_ARs. Spinophilin negatively regulates α_2_-adrenergic responses by blocking the association of G protein receptor kinase 2 with agonist-receptor-Gβγ complexes, thereby antagonizing β-arrestin-2-dependent receptor endocytosis (*12*). As spinophilin-null (*Spn*^***-***/-^) mice showed enhanced sensitivity to sedation elicited by α_2_-adrenergic stimulation (*12*), we wondered how α_2_-agonistic stimulation would affect the sedation response in *Sipa1l1*^-/-^ mice. In the rotarod assay, *Sipa1l1*^-/-^ mice were significantly more resistant to UK 14,304-evoked sedation than WT mice (Fig. 9A), suggesting a possible inhibitory role of SIPA1L1-spinophilin interactions in spinophilin-mediated repression of the α_2_-adrenergic response. To examine whether the resistance of *Sipa1l1*^-/-^ mice to sedation is generalized or non-specific in nature, we used another sedation-eliciting GPCR agonist, R-PIA, an agonist for adenosine A1 receptors. *Sipa1l1*^-/-^ mice unexpectedly showed an enhanced response to R-PIA-stimulated sedation (Fig. 9B), similar to that of *neurabin-1*^-/-^ mice (*17*). These results indicate a non-generalized, GPCR-pathway-dependent sedation response in *Sipa1l1*^-/-^ mice.

**Figure 9.**
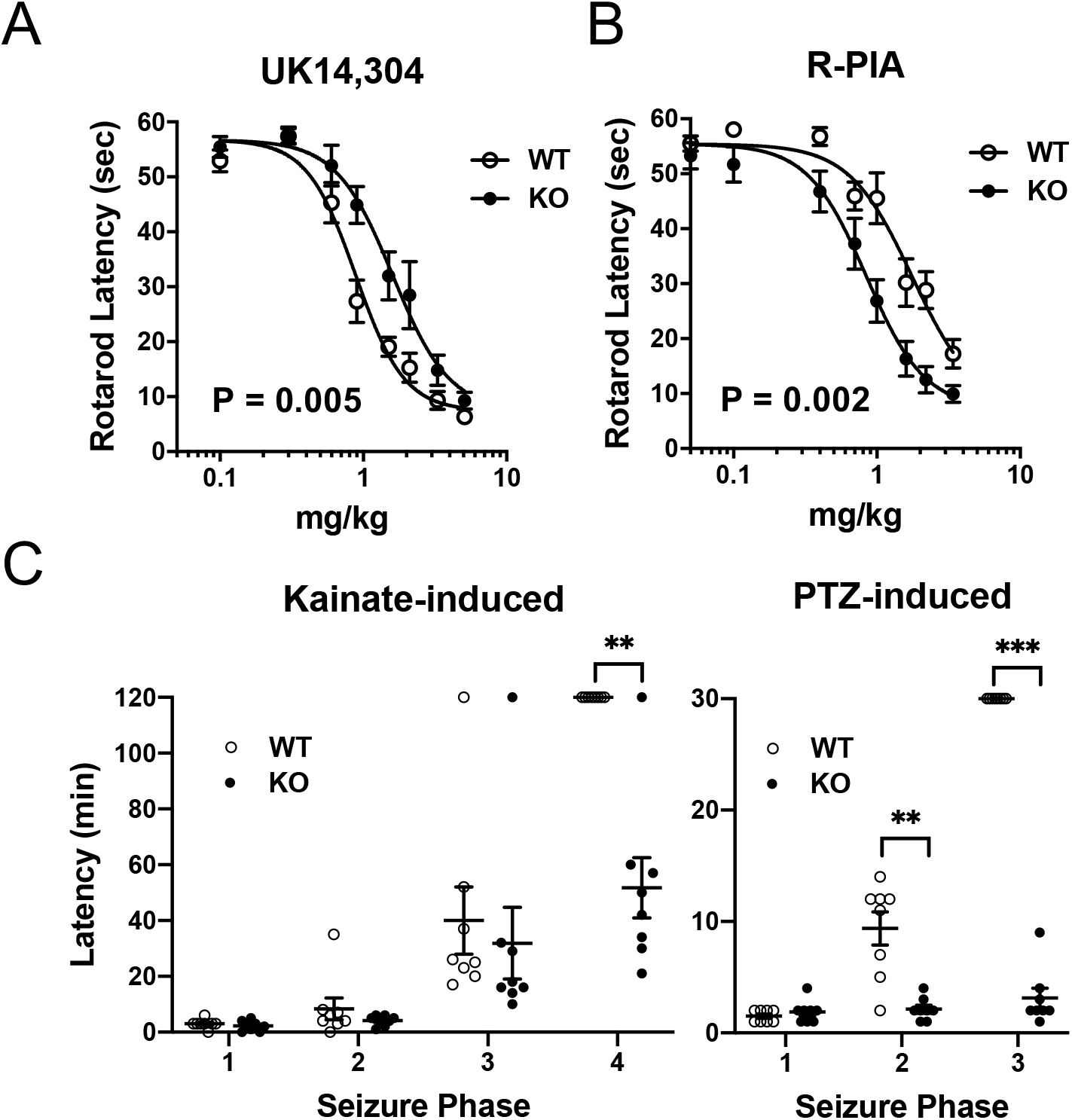
*Sipa1l1*^*-/-*^ mice show aberrant responses to GPCR agonist stimulation and significantly enhanced epileptic seizure susceptibility. (**A, B**) Sedation assessed by rotarod latency with increasing doses of α2AR agonist UK 14,304(A) or adenosine A1R agonist R-PIA (B). EC50 values for sedation in *Sipa1l1*^-/-^ (KO) and WT mice are (A) 1.60 and 0.89 mg/kg, (B) 0.84 and 1.76mg/kg, respectively. N = 11-12 per genotype. Values are mean ± SEM. P values indicate the genotype effect of two-way ANOVA. Partial η2 for genotype effect are 0.19 (A) and 0.34 (B). (**C**) Latency to manifest kainate- or PTZ-induced seizures. Phase 1, hypoactivity; phase 2, partial clonus; phase 3, generalized clonus; phase 4, severe generalized tonic-clonic seizure (see Methods for more details). Cutoff times are 120 or 30 min for kainate- or PTZ-induced seizures, respectively. No WT mice manifested phase 4 or 3 for kainate- or PTZ-induced seizures, respectively. N = 8 per genotype. Mean ± SEM is shown in the dot blots. **P < 0.01; ***P < 0.001. Detailed statistical information is available in Table S4.

Another interesting phenotype observed in *Spn*^***-***/-^ mice is their resistance to kainate-or pentylenetetrazole (PTZ)-induced seizures (*31*). Although the mechanism underlying this phenotype is not well understood, several lines of evidence show that the neurotransmitter norepinephrine and α_2_AR agonists exert powerful antiepileptogenic actions that are mediated by postsynaptic α_2A_ARs, one of the three α_2_AR subtypes (*32*). Moreover, α_2A_AR mutant mice show significantly enhanced epileptic seizure susceptibility (*33*). Thus, we hypothesized that *Sipa1l1*^-/-^ mice might also have enhanced seizure susceptibility. Indeed, *Sipa1l1*^-/-^ mice showed significantly enhanced susceptibility to kainate- or PTZ-induced seizures (Fig. 9C). An intraperitoneal injection of kainate (30 mg/kg) caused severe generalized tonic-clonic seizures (phase 4) in 7 out of 8 *Sipa1l1*^-/-^ mice, whereas no WT mice (0/8) reached phase 4. A subconvulsive injected dose of PTZ elicited no generalized clonus (phase 3) in WT mice (0/8), but all *Sipa1l1*^-/-^ mice (8/8) showed whole-body clonus with a sudden loss of upright posture.

### *Sipa1l1*^*-/-*^ mice show various types of behavioral impairment relevant to neuropsychiatric disorders

Since spinophilin is suggested to target various GPCRs, such as αARs, mAchRs, dopamine D2 receptors, μ-opioid receptors, and mGluRs, all of which are known to cause aberrant behaviors and to lead to neuropsychiatric disorders when dysregulated (see Discussion for details), we investigated consequences of the loss of SIPA1L1 through a series of behavioral tests.

*Sipa1l1*^-/-^ mice were born with the expected Mendelian ratio, were apparently healthy, and had lifespans similar to those of their WT littermates (773 ± 33 and 784 ± 36 days for WT and *Sipa111*^-/-^, respectively; mean ± S.E.M.; N=31 and 38 for WT and *Sipa1l1*^- /-^, respectively; U = 532.5, P = 0.50; two-tailed Mann-Whitney test). Gross anatomy of major organs, including brains of *Sipa1l1*^-/-^ mice, was comparable to that of WT littermates, and distribution and expression levels of major synaptic proteins were not affected in the *Sipa1l1*^-/-^ brain (Fig. S7). Nevertheless, *Sipa1l1*^-/-^ mice showed striking hyperactivity in the open field test (Fig. 10A, B), with significantly less time spent in the center area and more time spent close to the walls, which is considered an indication of increased anxiety (Fig. 10C, D). In the light-dark transition test, despite increased locomotor activity, *Sipa1l1*^-/-^ mice showed similar or slightly smaller transition numbers and significantly less time spent in the light chamber (Fig. 10E, F), also considered an indication of enhanced anxiety.

**Figure 10.**
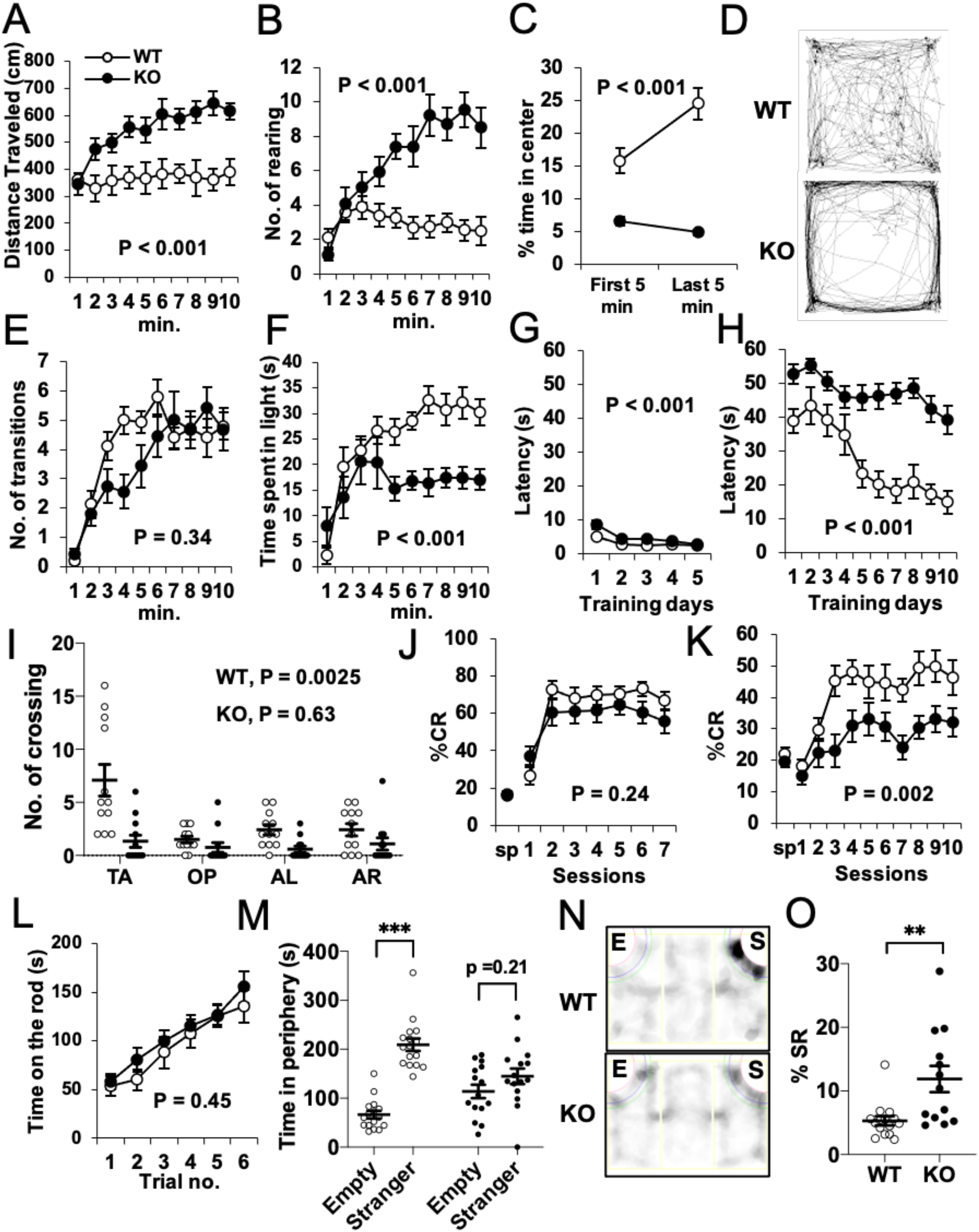
*Sipa1l1*^*-/-*^ mice show various types of behavioral impairment relevant to neuropsychiatric disorders. (**A**-**D**) Open field test. (D) Representative traces of mice in a 50 × 50 cm arena during a 10-min test. A 30 × 30 cm area in the center of the arena is defined as the center in (C). (**E, F**) Light-dark transition test. (**G**-**I**) Morris water maze. Control visible (G) and hidden (H) platform tests. Latency to reach the escape platform is shown. Mice were trained with 6 trials/day. In the probe test (I), the number of crossings of the original platform area or an equivalent area in each quadrant are shown. TA, target; OP, opposite; AL, adjacent left; AR, adjacent right. (**J, K**) Eyeblink conditioning. Results from delay (J) and trace (K) paradigms are shown. CR, conditioned response; sp, spontaneous eye blinking. (**L**) Accelerated rotarod test. (**M, N**) Three-chamber social interaction test. (M) shows total time spent in the peripheral areas of an empty cage or a cage with a stranger mouse. (N) Representative mountgraphs showing how long a mouse stayed in certain areas. E, empty cage; S, cage with stranger mouse. (**O**) The percentage of startle responses during the initial 30-ms period of the CS (conditioned stimulus) in eye blink conditioning. SR, startle response. Mean ± SEM is shown in line graphs and dot blots. **P < 0.01; ***P < 0.001. P values labelled in the line graphs indicate the genotype effect of two-way ANOVA. P values labelled in (I) indicate the quadrant effect of one-way ANOVA. N = 12-18. Detailed statistical information is available in Table S4.

As SIPA1L1 expression is enriched in the hippocampus, we tested hippocampus-dependent spatial learning by Morris water maze. Although *Sipa1l1*^-/-^ mice showed slight decrease in performance compared to WT mice in hippocampus-independent visible platform (control) test (Fig. 10G), the difference was minimal and *Sipa1l1*^-/-^ mice were able to reach similar level with WT mice by day five of the training (Table S4). However, in the hippocampus dependent hidden platform test (Fig. 10H) and subsequent probe test (Fig.10I), *Sipa11l*^-/-^ mice showed severely impaired learning even after 10 days of training. In the test of classical eyeblink conditioning, an associative learning that is not influenced by activity level (*34, 35*), *Sipa1l1*^-/-^ mice showed normal learning in the delay paradigm (Fig. 10J), which is dependent on cerebellar function, but impaired learning in the trace paradigm (Fig. 10K), which is a more complex learning task that depends on both the hippocampus and cerebellum. Regarding cerebellar function, *Sipa1l1*^-/-^ mice showed comparable motor coordination and learning with their WT littermates in the accelerating rotarod test (Fig.10L), suggesting that cerebellar function is not much affected in *Sipa1l1*^-/-^ mice.

In the three-chamber social interaction test, *Sipa1l1*^-/-^ mice manifested significantly reduced interest in stranger mice (Fig. 10M, N), suggesting autistic-like behavior. Recently, it has been shown that the acoustic startle eyeblink response is enhanced in patients with autism spectrum disorder (ASD) (*36, 37*). Enhanced acoustic startle response is also associated with Fragile X syndrome (FXS), the most prevalent cause of intellectual disability that is frequently accompanied by hyperactivity, autism, and/or seizures (*38*). We found that *Sipa1l1*^-/-^ mice show an enhanced acoustic startle eyeblink response (Fig. 10), similar to *Fmr1* mutant mice, a mouse model of FXS (*38*).

Collectively, these results demonstrate critical roles of SIPA1L1 in multiple behaviors that are relevant to neuropsychiatric disorders, such as attention deficit hyperactivity disorder (ADHD), anxiety disorder, intellectual disability, ASD, or FXS.

## Discussion

In this work, we have shown that, contrary to prevailing belief, SIPA1L1 is a not a major component of the PSD-95/NMDA-R complex, and that the vast majority of it does not localize to PSD. SIPA1L1 is suggested to be a cytoplasmic or submembranous protein distributed throughout neurons, interacting with the neurabin family of proteins, possibly to regulate GPCR signaling. *Sipa1l1*^-/-^ mice showed striking behavioral anomalies without obvious changes in spine size distribution or NMDA-R-dependent synaptic plasticity. On the other hand, *Sipa1l1*^-/-^ mice showed resistance or enhanced response to α_2_AR or adenosine A1 receptor agonist stimulation, respectively. These results demonstrated that SIPA1L1 deficiency could result in serious behavioral abnormalities that are relevant to neuropsychiatric disorders. This may open new avenues for research on disorders that involve spinophilin- or neurabin-1-regulated GPCR signaling.

The reasons for discrepancies between our work and previous studies are not all clear, but differences in materials and methods, e.g., primary cultured neurons vs neurons in mature brain or specificity of antibodies, may explain some of them. In addition, introduction of the cIP strategy, combined with stringent solubilization and wash conditions, could have made a difference in co-IP experiments in terms of minimizing artifactual interactions. This is particularly true for proteins that could physically bind each other *in vitro*, such as the case for SIPA1L1 and PSD-95 (*39*). Proteins that require a strong detergent for solubilization and subsequent neutralization for antibody binding would have a risk for artifactual interactions. We also speculate that since the actin cytoskeleton and its binding proteins are resistant to detergent solubilization (*40-42*), especially to non-ionic detergent such as Triton X-100, the non-PSD actin-binding proteins would be prone to contamination in conventional detergent extraction methods to determine PSD components. Thus, the pepsin pretreatment-immunostaining analysis could be a good alternative in determining the proportion of easy access (non-PSD) proteins and densely packed (PSD) proteins, as we have shown in this work. However, IEM will be the gold standard for conclusive results.

Frequent colocalization of SIPA1L1 and spinophilin throughout the cerebrum suggests that one of the major functions of SIPA1L1 involves interaction with spinophilin. This suggests an extrasynaptic and modulatory role, involving some GPCRs that are targets of spinophilin. In the case of α_2_AR signaling, the simplest model may be that SIPA1L1-spinophilin interaction inhibits the spinophilin-α_2_AR interaction, thus enhancing α_2_AR signaling. αAR signaling participates in multiple brain functions, including cognition. α_2_AR agonist stimulation could augment prefrontal cortex function, and is currently used to treat ADHD (*43*). One of the mechanisms underlying its efficacy may be targeting of hyperpolarization-activated cyclic nucleotide-gated (HCN) channels by α_2A_AR signaling, which localizes to the extrasynaptic region of dendritic spines (*44*). Furthermore, α_*1B*_*AR*^-/-^ mice showed hyperactivity and severely impaired learning in the Morris water maze (*45*). Thus, downregulation of αAR signaling may contribute to some of the behavioral anomalies in *Sipa1l1*^-/-^ mice.

*Sipa1l1*^-/-^ mice showed many characteristics common to FXS, which include hyperactivity, anxiety, intellectual disability, altered sensorimotor integration, autistic behavior, and susceptibility to seizures. A possible link between SIPA1L1 and FXS may be the regulation of Gp1 mGluRs via spinophilin (*16*). In the compelling “mGluR theory”, overactivation of mGluR function is postulated to mediate many symptoms of FXS, including learning deficits and seizure sensitivity (*46*). *Fmr1* mutant mice show enhanced mGluR-LTD, whereas *Spn*^-/-^ mice show decreased mGluR-LTD (*16*). Interestingly, *Sipa1l1* mRNA binds FMR1 (an RNA-binding protein that regulates translation) (*47*) and SIPA1L1 translation has shown to be up-regulated in juvenile, but significantly down-regulated in adulthood of *Fmr1* mutant mice (*48, 49*). Although most of the protein expressions were unchanged in adult *Fmr1* mutant brain, the expression of 14 proteins, including SIPA1L1, was significantly down-regulated to less than half compared to WT control (*49*). As treatment by mGluR antagonists could ameliorate phenotypes of adult *Fmr1* mutant mice (*50, 51*), down-regulation of SIPA1L1 may contribute to overactivation of mGluR function in mature *Fmr1* mutant mice and possibly in FXS patients, by enhancing spinophilin function. Alternatively, down-regulation of SIPA1L1 may simply contribute to behavioral anomalies of FXS in adulthood through other pathways. Whether SIPA1L1 has a role in regulating mGluRs and/or in FXS requires further study.

Dysregulation of other target GPCRs of spinophilin (μ-opioid receptors, mAchRs, and dopamine D2 receptors) or neurabin-1 (adenosine A1 receptors) may also contribute to some of the behavioral phenotypes in *Sipa1l1*^-/-^ mice. μ-opioid receptors are implicated in major depressive disorder (*52*), whereas mAchRs are involved in schizophrenia and Alzheimer’s disease (*53*). Dysregulation of the dopaminergic system has been implicated in a number of neuropsychiatric disorders and all currently available antipsychotics act via down-regulation of dopamine D2 signaling (*53*). D2 signaling was recently implicated in ASD and may promote social avoidance (*54*). Although regulation of dopamine D2 receptors by spinophilin is not well defined, *Spn*^-/-^ mice may have down-regulated D2 signaling (*55*). The SIPA1L1-neurabin-1 interaction could be related to the enhanced response of *Sipa1l1*^-/-^ mice to stimulation with adenosine A1 receptor agonists.

To our knowledge, genetic link between *SIPA1L1* and neuropsychiatric disorders is not clear to date, but it may be worth noting that a putative causal DNA variation of *SIPA1L1* in exome sequencing data of Australian ASD cofort has recently been reported (*56*). Further detailed study of molecular mechanisms involving the SIPA1L1-spinophilin (or neurabin-1) interaction and their target GPCR pathways will enhance understanding of mechanisms of higher brain functions and may provide novel perspectives in studies of neuropsychiatric disorders.

## Acknowledgements

We thank Hiroshi Toriyama, Toshihiro Maruyama and Miyuki Koumura at The University of Tokyo IMCB Olympus Bioimaging Center (TOBIC) for technical assistance with Olympus microscopes. We thank OIST Imaging Section and Paolo Barzaghi for their support with SD-OSR. We thank Masaki Sagara and Yoshihiro Kawasaki for providing plasmid constructs of spinophilin mutants. This work was supported by JSPS KAKENHI Grant Numbers JP24790311 and JP26460385.

## Author contributions

KM designed the study and performed most of the experiments and data analysis. S. Kobayashi and TM performed and analyzed electrophysiological experiments; K. Konno, MY, and MW performed and analyzed IEM. T. Horiuchi and S. Kawahara performed and analyzed eyeblink conditioning. TS performed electron microscopy. KI and RN performed mass spectrometry. T. Hayashi, K. Kohu, and NY-S generated anti-SIPA1L family antibodies. KM and KS generated *Sipa1l1*^-/-^ mice. TN, TY, YK and TA supervised the study. KM and TA wrote the paper.

## Data Availability

Most of the data generated or analyzed during this study are included in this article and its supplementary information files. Other datasets generated during and/or analysed during the current study are available from the corresponding authors on reasonable request.

## Methods

### Antibodies

Polyclonal anti-SIPA1L1, SIPA1L2, and SIPA1L3 antibodies were prepared by immunizing rabbits with fragments of SIPA1L1 (amino acids 1617 – 1804 of accession no. NP_056371.1), SIPA1L2 (amino acids 1 – 70 of accession no. NP_065859.3) or SIPA1L3 (amino acids 1049 – 1251 of accession no. NP_055888.1) fused to glutathione S-transferase. Antibodies were purified with affinity chromatography using columns to which antigens used for immunization had been linked. Other antibodies used in this work are listed in Table S5.

### Targeted disruption of the Sipa1l1 gene

Genomic clones for the *Sipa1l1* locus were isolated by screening with a C57BL/6N male liver genomic library (Clontech). The targeting vector was constructed by inserting a nuclear localization signal-*LacZ*-polyA cassette, followed by a *PGK*-*neo*-polyA cassette at the first methionine site, preserving 0.8 kb (5’) and 5.2 kb (3’) of the flanking regions (Fig S1A). TT2 ES cells were electroporated and selected by standard procedures. Correctly targeted clones were screened by PCR and subsequently confirmed by Southern blotting. Targeted clones were used for aggregation with eight-cell embryos, and chimeric males were mated with C57BL/6N females. Subsequent genotyping was performed by genomic PCR. Primers used were FW: 5’-TAGATCCGTGTGCCACAA-3’, RV: 5’-GAGGCCAATCTGCTATTC-3’, and LacZ: 5’-CAGTCACGACGTTGTAAAAC-3’. Heterozygotes were then backcrossed to C57BL/6N mice for at least 9 generations. 2-to 4-month-old male mice were used for all the experiments. They were kept in a 14-h light/10-h dark cycle in a temperature- and humidity-controlled (22-24°C, 50-60%) specific pathogen-free vivarium, and they had *ad libitum* access to food and water. All animal experiments were conducted according to guidelines for care and use of animals, approved by the Animal Experiment Committee of Institute for Quantitative Biosciences, The University of Tokyo.

### Immunohistochemistry and X-Gal staining

Mice were deeply anesthetized with 90 mg/kg sodium pentobarbital and were intracardially perfused with ice-cold sodium phosphate buffer (pH 7.3, NPB), followed by ice-cold 4% paraformaldehyde (PFA) /NPB. The whole brain was removed, separated bilaterally at the medial line and fixed in ice-cold 4% PFA/NPB for 2 h. The brain was further infiltrated sequentially with 10, 15, and 20% sucrose/NPB for more than 4 h at each concentration and then frozen in a Tissue-Tek OCT compound (Sakura Finetek). 10-μm cryosections were attached to an MAS-coated slide glass (S9441 Matsunami) and air dried for 2 h. For permeabilization, sections were incubated in 0.3% Triton X-100/Tris-buffered saline (pH7.5, TBS) for 10 min at room temperature (RT) except for synaptophysin staining, which sections were incubated in boiling 10 mM sodium citrate for 5 min. Alternatively, they were incubated in 0.4 mg/mL pepsin in 0.2 N HCl for 2 min at 37°C for a PSD localization assay. Sections were blocked with TBS containing a 0.5% blocking reagent (Roche), 2% fetal bovine serum, and 0.1% Tween-20 for 1 h. Then they were incubated overnight at 4°C with primary antibodies diluted in the blocking buffer. In the case of reactions containing mouse antibodies, reagents from the VECTOR M.O.M. Basic Kit (Vector Laboratories) were added. Following washes in TBS containing 0.1% Tween-20 (TBST), sections were incubated for 1 h at RT with secondary antibodies diluted in blocking buffer. Sections were subsequently stained with TOPRO-3 or DAPI, washed, and coverslipped with Vectashield mounting medium (Vector Laboratories). Sections from WT and KO mice were processed simultaneously on the same slide glass. Antibodies and their dilutions used for immunostaining are listed in Table S5. A Zenon Rabbit IgG Labeling Kit (Molecular Probes) was used for staining neurabin-1 according to manufacturer’s instruction. Briefly, labelling of SIPA1L1 was performed by regular protocol as above to secondary antibody incubation. After TBST washes, sections were incubated with Zenon-labelled (molar ratio 6:1) neurabin-1 antibody for 1.5 h at RT. After three TBST and two PBS washes, sections were fixed with 4% PFA for 15 min, washed, and coverslipped with Vectashield mounting medium. Parallel experiment using negative-control omitting neurabin-1 antibody in Zenon labelling reaction resulted in no significant signal. For diaminobenzidine (DAB) staining, Vectastain Elite ABC and Avidin/Biotin Blocking Kits (Vector Laboratories) were used in conjunction with the above procedures. For X-Gal staining, dried sections were stained overnight at 37°C in an X-Gal staining solution and subsequently counterstained with Nuclear Fast Red (Vector Laboratories). Nissl staining was performed by incubating sections in a 1% thionin solution for 60 min at RT. Digital images were obtained using commercial Olympus microscopes. Briefly, low magnification images were obtained by IX-83 (Olympus) equipped with DP80 camera and motorized stage using x 20 objective. Whole brain section or region of interest was scanned and stitched automatically using cellSense software (Olympus). For confocal imaging, FV1000 (Olympus) was used with x 100, 1.4 NA silicone immersion objective UPLSAPO100XO. For SRM, SD-OSR (Olympus) equipped with Yokogawa CSU-W1 scanner, Hamamatsu Orca Flash 4 V2+ High Speed SCMOS camera, and x 100, 1.35 NA silicone immersion objective with correction collar (UPLSAPO100XS) was used to acquire Z-stack images (200-nm step size, all channels scanned in each plane) of 1024 × 1024 pixels (41 × 41 μm^2^)/image. Original images adjusted only for brightness and contrast by Fiji/ImageJ (NIH) are shown in the figures.

### Immunoelectron microscopy

Mice were fixed by transcardial perfusion with 3% glyoxal-based fixative (*57*). Brains were cryoprotected with 30% sucrose in 0.1 M phosphate buffer to prepare 50 μm-thick cryosections on a cryostat (CM1900; Leica Microsystems). All immunohistochemical incubations were performed at room temperature. For silver-enhanced pre-embedding immunogold electron microscopy, microslicer sections were dipped in 10% normal goat serum / PBS for 30 mins, incubated overnight with SIPA1L1 antibody (1:1000) diluted with 0.1% TritonX-100 /PBS, and subjected to silver-enhanced immunogold labeling using anti-rabbit IgG conjugated with 1.4 nm gold particles (Nanogold; Nanoprobes, USA) and R-Gent SE-EM Silver Enhancement Reagents (Aurion, Netherlands). Sections were further treated with 1% osmium tetroxide and 2% uranyl acetate, and embedded in Epon812. Ultrathin sections (100 nm in thickness) were prepared with an ultramicrotome (Leica, Wien, Austria), and photographs were taken with an H7100 electron microscope (Hitachi, Tokyo, Japan). The density and distribution of immunogold particles were quantitatively analyzed on electron micrographs using MetaMorph software (Molecular Devices; n = 2 mice). The density of SIPA1L1 on excitatory nerve terminals and dendritic spines was calculated by measuring the number of immunogold particles. Perpendicular distribution of PSD-95 or SIPA1L1 was examined by sampling synaptic profiles whose presynaptic and postsynaptic membranes were cut perpendicularly to the plane of the synaptic cleft, and by measuring the distance from the midline of the synaptic cleft to the center of immunogold particles. Statistical significance was assessed by one-way analysis of variance (ANOVA) with Tukey’s multiple comparison and Kolmogorov–Smirnov test.

### Spine size analysis by electron microscopy

Electron microscopy was basically performed as described (*21*). Briefly, littermate mice (2-3 months) were deeply anesthetized with sodium pentobarbital and were intracardially perfused for 5 min with 2.5% glutaraldehyde in a 0.1 M phosphate buffer (pH 7.4).

Hippocampi were removed from whole brains, and the CA1 areas of hippocampi were cut into tiny blocks. These blocks were postfixed in the same fixative for 3 h, osmicated with 1% osmium tetroxide in a 0.1 M phosphate buffer for 2 h, washed thoroughly with a 5% sucrose solution, dehydrated in a graded alcohol series, embedded in Epok812 (#02-1001, Okenshoji Co.), and cured for 12 h at 60°C. For each block, 1-μm sections were cut and stained with 1% toluidine blue to guide further trimming to isolate the equivalent CA1 subfields. Ultra-thin sections (80 nm) were cut with a diamond knife and stained with uranyl acetate and lead citrate, and then observed with a JEOL JEM1010 electron microscope operated at 100 kV. Similar neuropil areas of stratum radiatum not containing cell bodies or blood vessels were randomly selected within 100–250 μm of the CA1 pyramidal cell body and photomicrographed at 5,000× magnification. Five electron micrographs representing 1300-1400-μm^2^ neuropil regions in each mouse were taken. Image negatives were scanned at 1200 dpi and analyzed with ImageJ. The number of synapses (synapse density), PSD lengths, and cross-sectional areas of spine heads from 4 mice per genotype were quantified. Excitatory synapses bearing spines were defined by the presence of a clear PSD facing at least three presynaptic vesicles. Measurements were performed by an experimenter blind to the genotype.

### Electrophysiological analysis

Standard procedures and solutions described previously (*58*) were used. In brief, hippocampal slices (400 µm) were prepared from mice 8-12 weeks of age. Synaptic responses were recorded at 25.0 ± 0.5°C with extracellular field-potential recordings in the stratum radiatum of the CA1 region using a glass recording pipette filled with 3 M NaCl. External solution contained the following (in mM): 119 NaCl, 2.5 KCl, 1.3 MgSO_4_, 2.5 CaCl_2_, 1.0 NaH_2_PO_4_, 26.2 NaHCO_3_, 11 glucose, and 0.1 picrotoxin (a GABA_A_-receptor antagonist). To evoke synaptic responses, Schaffer collateral/commissural fibers were stimulated at 0.1 Hz (test pulse) with a bipolar tungsten electrode. Stimulus strength was adjusted to evoke excitatory postsynaptic potentials (EPSPs) with a slope of 0.10-0.15 mV/ms, except for the experiments examining input-output relationships. Input–output relationships were examined in the presence of a low concentration of the non-NMDA receptor antagonist, 6-cyano-7-nitroquinoxaline-2,3-dione (CNQX: 1 μM) in the external solution to partially block AMPA receptor-mediated EPSPs. This partial blockade enables more accurate measurements since the nonlinear summation of field EPSPs is reduced. EPSPs were evoked with various strengths of stimulation, and data were first sorted by binning fiber volley amplitudes. Then EPSP amplitudes were averaged within each bin. The paired-pulse facilitation was examined in the presence of 25 µM D-(–)-2-amino-5-phosphonopentanoic acid (D-AP5). Paired-pulse stimuli at intervals of 50, 100, and 200 ms were applied every 10 s. An Axopatch-1D amplifier (Molecular Devices, Sunnyvale, CA, USA) was used to record EPSPs. Data were digitized at 10 kHz and analyzed on-line using pClamp software (Molecular Devices). All values were reported as means ± standard errors of the mean. Student’s t-test was used to determine whether there was a significant difference in the means of two datasets. Picrotoxin was purchased from Sigma-Aldrich (St. Louis, MO, USA). D-AP5 and CNQX were purchased from Tocris Bioscience (Avonmouth, UK).

### Colocalization analysis

Colocalization analysis was performed using GDSC ImageJ plug-in according to developer’s Colocalization User Manual (http://www.sussex.ac.uk/gdsc/intranet/microscopy/UserSupport/AnalysisProtocol/imagej/colocalisation). Briefly, SRM images stained for SIPA1L1 and its candidate interacting-proteins at the neuropil region of layer V cerebral cortex or hippocampal CA1 area were acquired using Olympus SD-OSR as described above. Four serial Z-stack images (200-nm step size) of 1024 × 1024 pixels (41 × 41 μm^2^)/image were processed to define foreground and background by Otsu method (or Triangle method for images with relatively low signal-to-noise ratio) using Stack Threshold Plugin. The processed images and original images were used to calculate statistical significance of Manders coefficient and Pearson correlation coefficient, respectively, by Confined Displacement Algorithm Plugin. If the presence of irregular structures such as somata interfered with the analysis, corresponding region was excluded from the confined region. Random displacement was defined using radial displacement chart for Pearson correlation coefficient for each sample. A *P* value of < 0.01 was adopted for statistically significance.

### Cell culture, transfections, immunostaining, and immunoprecipitation

HEK293T or COS-7 cells were cultured in Dulbecco’s modified Eagle’s medium supplemented with 10% fetal bovine serum at 37°C, 5% CO_2_. Expression vectors for the SIPA1L family of proteins were generated by cloning human *SIPA1L1* (NM_015556.3), *SIPA1L2* (AY168879), *or SIPA1L3* (AY168880) cDNA into FLAG- or Myc-pcDNA3.1 (+). Expression vectors for spinophilin have been described elsewhere (*59*). Transfections of plasmid constructs were performed using Lipofectamine 2000 (Invitrogen) for HEK293T and Lipofectamine LTX (Invitrogen) or TransIT-LT1 (Mirus Bio) for COS-7 cells, according to manufacturers’ instructions. Immunostaining was performed as described previously (*59*). For immunoprecipitation, COS-7 cells were lysed in lysis buffer (0.33 % SDS, 1.67 % Triton X-100, 50 mM Tris-HCl pH7.4, 150 mM NaCl, 1 mM EDTA, 1 mM EGTA, 1mM PMSF, 1mM Na_3_VO_4_, 25mM NaF, 5 µg/mL aprotinin, chymostatin, leupeptin and pepstatin A, 10% glycerol), rotated for 60 min at 4°C and centrifuged at 17,000 × g for 40 min. Supernatants were pre-cleared with Dynabeads Protein G (Invitrogen Dynal) and incubated with anti-Myc (MBL) antibody-bound Dynabeads Protein G with overnight rotation at 4°C. Samples were then washed 4 times with wash buffer (0.33% SDS, 1.67%Triton X-100, 50 mM Tris-HCl pH7.4, 150 mM NaCl, 1 mM EDTA, 1 mM EGTA, 10%glycerol). Proteins were eluted by incubation in 50 mM Tris, pH 6.8 with 2% SDS for 10 min at room temperature with shaking. Samples were analyzed by Western blotting, as above.

### cIP-MS and cIP-WB analyses

Mouse brain regions of interest were quickly dissected on ice-cold filter paper soaked with homogenization buffer (HB; 0.32M Sucrose, 20 mM HEPES pH7.4, 1 mM EDTA, 1 mM EGTA, 5mM NaF, 1mM Na_3_VO_4_) and were homogenized in ice-cold HB using a Dounce homogenizer. After homogenates were centrifuged at 800g for 10 min, supernatants were transferred to new tubes, and proteins were crosslinked by adding 20 mM DSP (dithiobis [succinimidylpropionate], Thermo Scientific Pierce), a primary amine-reactive and membrane-permeable crosslinker with a 1.2-nm spacer arm, to a final concentration of 200 µM. For non-crosslinked controls, the same volume of solvent (DMSO) was added. Tubes were rotated at 4°C for 10 min, and 1 M Tris-Cl (pH 7.4) was added to a final concentration of 100 mM to terminate the crosslinking reaction. After 15 min of rotation, tubes were centrifuged at 9,200 × g to obtain the P2 fractions, containing crude synaptosomes and plasma membranes. P2 pellets were solubilized in lysis buffer (2% SDS, 50 mM Tris-HCl pH7.4, 150 mM NaCl, 1 mM EDTA, 1 mM EGTA, 1mM PMSF, 1mM Na_3_VO_4_, 25mM NaF, 5 µg/mL aprotinin, chymostatin, leupeptin and pepstatin A, 10% glycerol) at 37°C for 30 min. Five times the volume of ice-cold neutralization buffer (2% Triton X-100, 50 mM Tris-HCl pH7.4, 150 mM NaCl, 1 mM EDTA, 1 mM EGTA, 1mM PMSF, 1mM Na_3_VO_4_, 25mM NaF, 5 µg/mL aprotinin, chymostatin, leupeptin and pepstatin A, 10% glycerol) was added and centrifuged at 17,000 × g for 1 h. Supernatants were pre-cleared using Dynabeads Protein G (Invitrogen Dynal) and incubated with anti-SIPA1L1, anti-PSD-95 (UPSTATE), or control IgG antibody with overnight rotation at 4°C. Samples were then rotated with Dynabeads Protein G for 90 min and washed 4 times with wash buffer (0.33% SDS, 1.67% Triton X-100, 50 mM Tris-HCl pH7.4, 150 mM NaCl, 1 mM EDTA, 1 mM EGTA, 10% glycerol). Proteins were decrosslinked and eluted by incubation in a 2x SDS sample buffer containing 200 mM DTT for 60 min at 37°C followed by 10 min at 56°C with constant mixing at 1400 rpm on an Eppendorf ThermoMixer. Dynabeads and unsolubilized materials were carefully removed magnetically and by centrifugation. Final supernatants were analyzed by SDS-PAGE.

For silver staining analysis, Perfect NT Gel (5-20%, DRC Co.) was used for SDS-PAGE. Staining and destaining were performed using SilverQuest (Invitrogen), according to the manufacturer’s protocol. Selected bands or corresponding areas in control lanes were excised and cut into 1 mm cubes, destained, and reduced with DTT (10 mM in 100 mM NH_4_HCO_3_, 56°C for 60 min) followed by alkylation with iodoacetamide (55 mM in 100 mM NH_4_HCO_3_ RT for 45 min). After repeated alternate washings with 100 mM NH_4_HCO_3_ and acetonitrile, gel pieces were rehydrated with 10 µL 50 mM NH_4_HCO_3_ containing 25 µg/mL trypsin (Trypsin Gold, Promega) and incubated for 15 min on ice. 10 µL of 50 mM NH_4_HCO_3_ was added, and trypsin digestion was carried out overnight at 37°C. Peptides were extracted with 20 µL of 20 mM NH_4_HCO_3_, followed by 20 µL of 50% acetonitrile, 0.1% trifluoroacetic acid (TFA) three times. The volume of pooled supernatants was reduced to 10-20 µL by vacuum centrifugation and then loaded into an automated electrospray ionization (ESI)-MS/MS system, which consisted of the DiNa system (KYA Tech Corporation) equipped with a C-18 ESI capillary column (100 μm × 150 mm, NIKKYO Technos) and an LTQ Velos Orbitrap ETD instrument (ThermoFischer Scientific). For protein identification, spectra were processed using Proteome Discoverer Version 1.2 (ThermoFisher Scientific) against SEQUEST with a 5% false discovery rate (FDR) cutoff. Experiments were performed one time each for cerebral cortex and hippocampus. The candidate SIPA1L1-interacting proteins were defined as proteins detected only in WT samples (PSMs ≧ 2)or PSMs of WT is more than 10-fold of that of KO samples.

Western blotting was performed by standard methods. Briefly, proteins were transferred to PVDF membranes (Immobilon, Millipore) and 5% skim milk, 0.1% Tween-20 in TBS was used for blocking and antibody dilution. Antibodies and associated dilution factors are listed in Table S5. A chemiluminescent signal was detected using Luminata Forte Western HRP (Millipore) and ImageQuant LAS4000mini (FujiFilm)

### GPCR agonist stimulation (sedation) analysis

9 to 10-week-old mice were evaluated in the rotarod test for sedation, basically as described previously (*12*). Briefly, the subject was placed on a rotarod (O’hara & Co.) rotating constantly at 10 rpm. Mice were trained for 3-6 sessions until they learned to remain on the rod for 60 s. Mice were injected *i*.*p*. with increasing doses of the α_2_AR-agonist, UK 14,304 (abcam), or the adenosine analog, (-)-N6-(2-Phenylisopropyl) adenosine (R-PIA; Sigma Aldrich) dissolved in saline. 10 min post-injection, each mouse was tested three times in succession for its ability to remain on the rotarod. Results of the three trials were averaged. Cumulative doses of agonists are shown in Fig. 9. The cutoff time was 60 s. The experimenter was blinded to the genotype during testing.

### Seizure susceptibility analysis

8 to 9-week-old mice were injected *i*.*p*. with 30 mg/kg of kainate (Sigma Aldrich) or PTZ (Sigma Aldrich) dissolved in saline in a volume of 15 mL/kg. Mice were placed in a clear Plexiglas cage and video-recorded for up to 2 h or the 30 min cutoff time for kainate- or PTZ-induced seizures, respectively. Seizures were scored according to the following scale. Phase 1, hypoactivity: behavioral arrest for more than 10 s; phase 2, partial clonus: a brief seizure, typically lasting 1 or 2 s, with clonic seizure activity affecting the face, head or forelimbs; phase 3, generalized clonus: the sudden loss of upright posture, whole body clonus involving all four limbs and tail, typically lasting for 30-60 s, followed by a quiescent period; and phase 4, severe generalized tonic-clonic seizure: a continuous loss of upright posture, lying or rolling on the floor, resulting in death from continuous convulsions. The experimenter was blinded to the genotype during testing.

### Behavioral analysis

Male *Sipa1l1*^-/-^ and WT mice were housed together, with two to four littermates (or mice with close birthdays) per cage after weaning. Mice were acclimated to handling and the experimental room for at least three days before the start of an experiment. An independent group of mice (2-3 months) was used for each test unless otherwise noted. Experimenters were blinded to the genotype during testing. All experiments were analyzed using an automated system from O’hara & Co., except during eyeblink conditioning. All Image series software (O’hara & Co.) used for analysis is based on the public domain NIH Image or ImageJ program (https://imagej.nih.gov/nih-image/).

### Open field test

Each subject was placed in the center of an open-field apparatus (50 × 50 × 33.3 cm; W × D × H) illuminated at 20 lux and allowed to move freely for 10 min. Distance travelled in the arena, trace of the movement, rearing activity, and time spent in the center were recorded and analyzed using Image OF 2.15x and Image OFC 2.03sx. Rearing activity was counted manually using the human observation mode of Image OFC 2.03sx. Accelerating rotarod and contextual and cued fear conditioning tests were subsequently performed on the same group of mice with an interval of two days between tests.

### Accelerating rotarod test

Mice were placed on a rod (3 cm in diameter) rotating at 4 rpm initially, and then the rotation of the rotarod was accelerated linearly to 40 rpm over a 300-s period. The latency to fall off the rotarod during a trial was automatically measured. Mice were trained for two consecutive days, receiving three trials per day at intervals of 90 min between trials.

### Light-dark transition test

The apparatus consisted of a box (21 × 42 × 25 cm) divided into two sections of equal size by a partition with a door. One chamber was brightly illuminated (100 lux), whereas the other chamber was dark without illumination. Mice were placed on the dark side and allowed to move freely between the two chambers with the door open for 10 min. The total number of transitions, time spent on each side, latency to enter the light side, and distance traveled were recorded and analyzed automatically using ImageJ LD1.

### Morris water maze

A pool with a 1-m diameter was filled with opaque water colored with nontoxic white paint and maintained at approximately 25°C. Each training trial began by placing the mouse in the quadrant that was either right, left, or opposite to the target quadrant containing a submerged platform (10-cm diameter), in semi-random order. The same order of start positions was used for all subjects. Training trials were a maximum 60 s in duration. A mouse that failed to reach the platform within 60 s was subsequently guided to the platform. Mice that reached or were guided to the platform stayed there for 20 s. Two trials per block with a 1-min inter-trial interval, three blocks per day with a 1-h inter-block interval were conducted for 10 or 5 days to train mice for hidden or visible platform tasks, respectively. The visible platform test was conducted after the completion of the hidden platform test. Latency to reach the platform, distance traveled to the platform, and average swim speed were automatically recorded. At the end of the tenth day of hidden platform training, a probe test was conducted for 1 min to confirm that spatial learning had been acquired, based on navigation by distal environmental room cues. Time spent in each quadrant and the number of crossings above the original platform site were automatically recorded. Data were automatically analyzed using Image WM 2.12r, Image WMV 2.08sr, and Image WMH 2.08s.

### Three-chambered social interaction test

4-month-old mice were used for this social interaction test. The testing apparatus consisted of a rectangular, three-chambered box and a lid with an LED light panel and a CCD monochrome camera. Each chamber was 20 × 40 × 22 cm, and separating walls were made from transparent Plexiglas with small openings (5 × 3 × 3 cm). The subject mouse was first placed in the middle chamber and allowed to habituate to the entire test box for 10 min. After habituation, the mouse was taken out of the box, and an age-matched unfamiliar WT male (stranger mouse), which had no prior contact with the subject mouse, was placed in a small, round wire cage in one of the side chambers. The side on which the stranger mouse was placed was systematically alternated between trials. The subject was placed back in the central chamber for a 10-min session, and the time spent in each chamber, the number of entrances to each chamber, the distance traveled in each chamber or in the periphery of each cage, the total distance travelled, the average travel speed, and the mountgraphs were automatically recorded and analyzed using TimeCS1 software (O’hara & Co.).

### Eyeblink conditioning

Mice were prepared for eyeblink conditioning basically according to previously described procedures (*34*). In brief, under anesthesia with pentobarbital and, if necessary, with diethyl ether inhalation, four Teflon-coated stainless-steel wires (No. 7910, A-M Systems) were implanted under the left upper eyelid. Two of these wires were used to record the eyelid electromyograms (EMGs), and the remaining two delivered an unconditioned stimulus (US). 2-3 days after surgery, mice were subjected to two days of habituation without US or conditioned stimulus (CS), during which EMGs were recorded to calculate spontaneous eyeblink frequency. 7 or 10 days of delay or trace conditioning, respectively, began the next day. A daily session consisted of 90 CS-US paired trials and 10 CS-alone trials at every 10^th^ trial. The CS was a 350-ms tone (1 kHz, 90 dB) with a 5-ms rise and a 5-ms fall time. The US was a 100-ms periorbital shock (100 Hz square pulses) with the intensity carefully adjusted to elicit a head-jerk response in each animal. The interstimulus interval was 250 ms or 850 ms in delay or trace conditioning, respectively. Eyelid EMGs were analyzed as described previously (*34*), except that trials that elicited a startle response to the CS were also included for evaluation of conditioned response (CR) occurrence. In brief, the mean + S.D. of amplitudes of EMG activity for 300 ms before CS onset in 100 trials was defined as the threshold, which was then used in the analysis below. In each trial, average values of EMG amplitude above the threshold were calculated for 300 ms before CS onset (pre-value), 30 ms after CS onset (startle-value), and 200 ms before US onset (CR-value). If the pre-value was < 10% of threshold, the trial was regarded as valid. Among valid trials, a trial was assumed to contain the CR if the CR value was larger than 1% of the threshold and it exceeded two times the pre-value. For the CS-alone trials, the period for CR-value calculation was extended to the CS end. To evaluate the effect on the startle response, we calculated the frequency of trials in which the startle-value exceeded 10% of the threshold.

### Statistical analysis

All results are expressed as means ± SEM, unless noted otherwise. Statistical analyses in this work employed unpaired two-tailed Student’s *t* tests, two-tailed Welch’s *t* tests, two-tailed Mann-Whitney tests, two-tailed Wilcoxon matched pair signed-rank tests, Kolmogorov-Smirnov tests, one-way ANOVA with Geisser-Greenhouse correction followed by Tukey’s post-hoc tests, or a two-way ANOVA with Geisser-Greenhouse correction followed by Sidak’s post-hoc tests, where appropriate using GraphPad Prism 8. A *P* value of < 0.05 was considered statistically significant. For effect size calculations, Pearson’s r or partial η^2^ was used. D’Agostino-Pearson test and F-tests were used to check normality and equal variance, respectively. Detailed statistical information is available in Table S4.

### Availability of materials

Most materials are readily available from commercial sources or from our lab. Exceptions are the rabbit polyclonal anti-SIPA1L1, 2, or 3 antibodies that we generated, or antibodies discontinued from commercial suppliers, due to limited amount. However, they may be available for reasonable requests.

---

**Figure S1.**
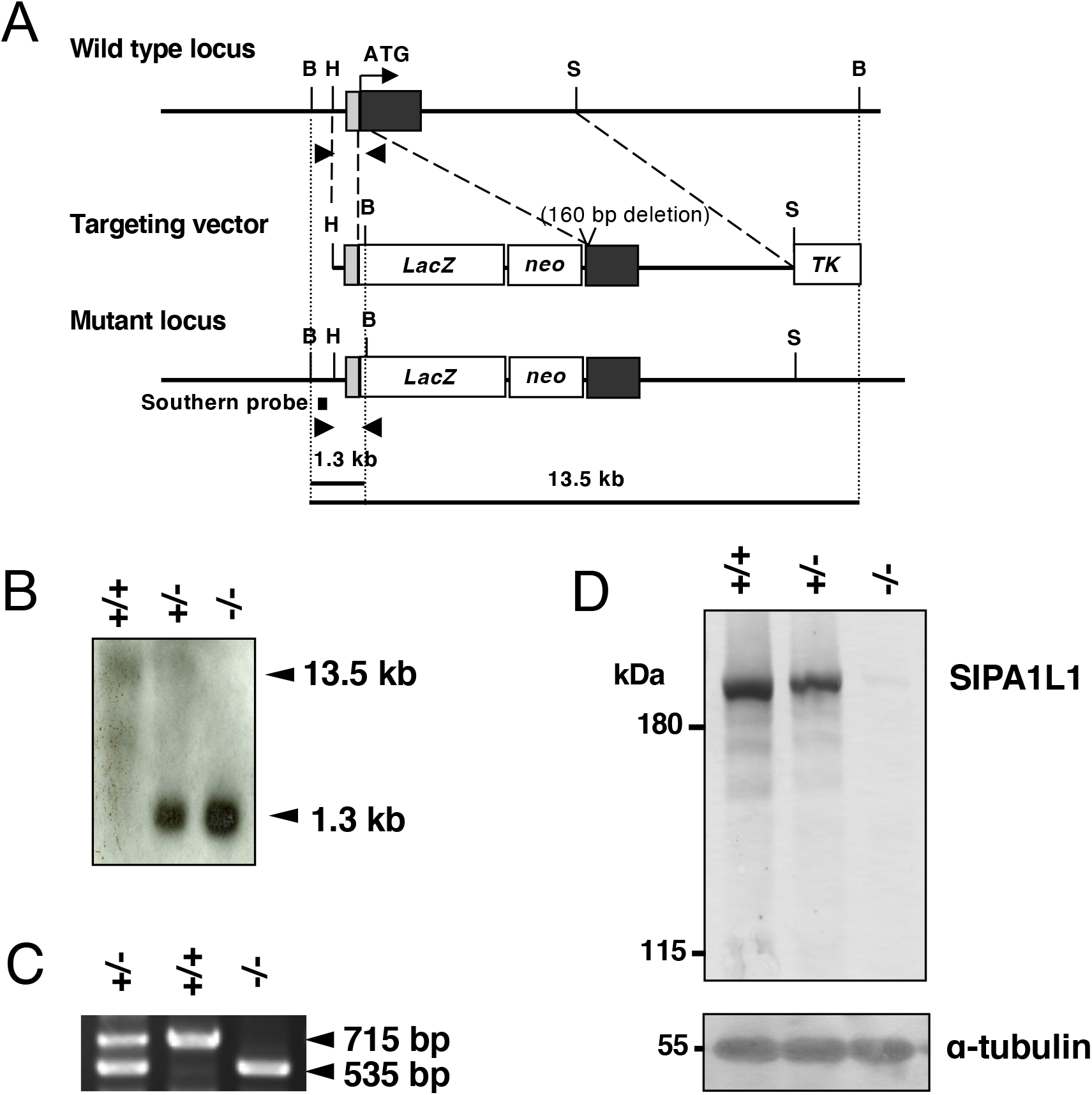
Targeted disruption of the *Sipa1l1* gene. **(A)** restriction map. Coding and noncoding regions in exons are indicated by black and grey boxes respectively. Arrowheads indicate primer positions used in PCR genotyping. B, BamHI; H, HindIII; S, SphI; neo, neomycin resistance gene; TK, herpes simplex virus thymidine kinase. (**B**) Southern blot analysis of genomic DNA extracted from mouse tail. Genomic DNA was digested with BamH I. The expected 13.5-kb (wild-type locus) and 1.3-kb (targeted locus) fragments were generated. (**C**) Genotyping of progeny of *Sipa1l1*^+/-^ intercrosses by PCR. (**D**) Western blot analysis of the brain lysate from mature mice.

**Figure S2.**
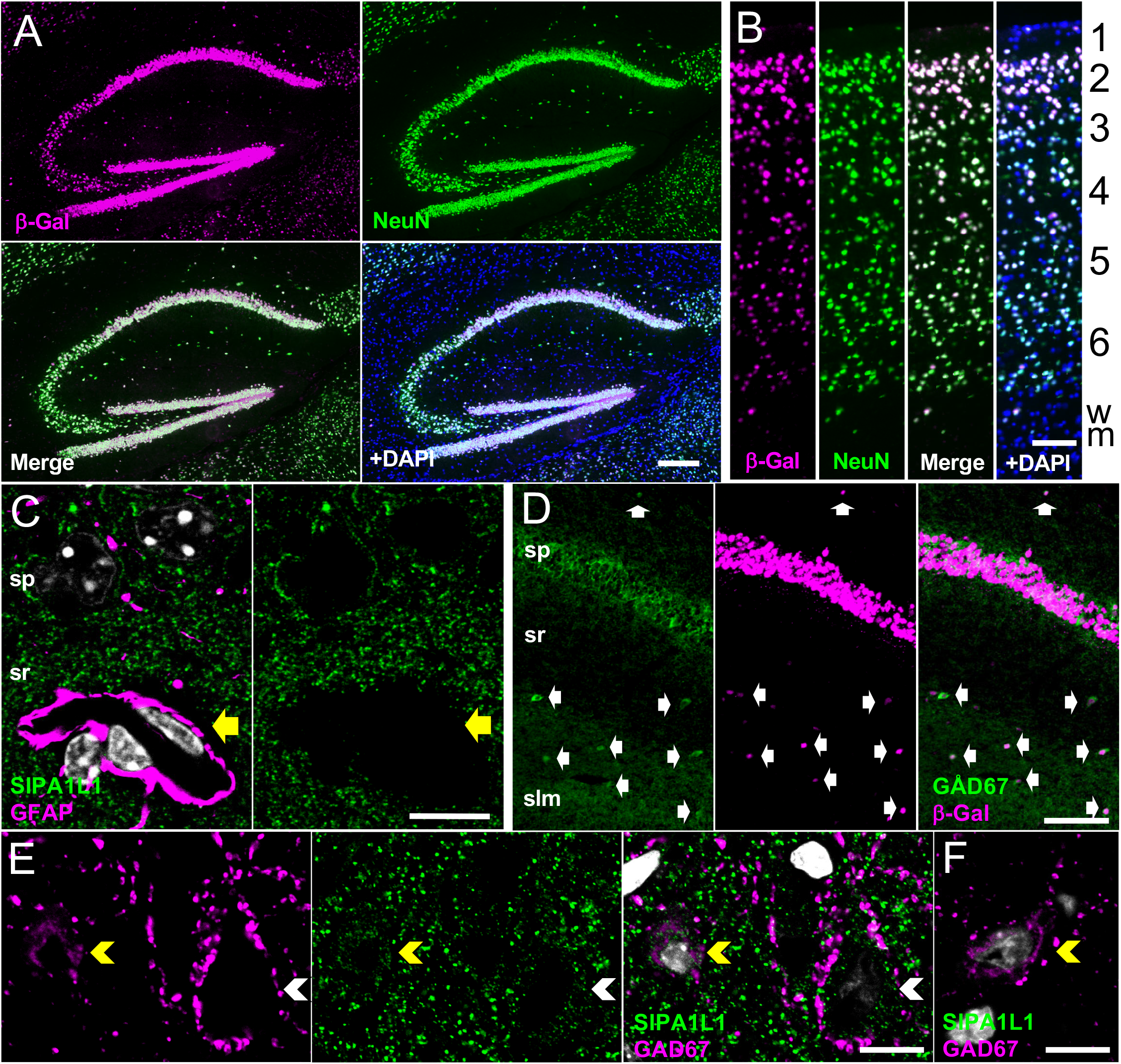
SIPA1L1 is expressed in excitatory and inhibitory neurons but not in glial cells. (**A, B**) Co-staining of β-Galactosidase (β-Gal) and NeuN in mature *Sipa1l1*^*+/-*^mouse brain. (A) and (B) show hippocampus and visual cortex, respectively. Nuclear DAPI staining is added in the fourth panels to show the whole population of cells. The numbers on the right indicate layers of visual cortex. wm, white matter. (**C**) GFAP, SIPA1L1, and nuclei (grey) in the hippocampal CA1 region. SIPA1L1 is not detected in GFAP-positive glial cells, indicated by the yellow arrow. (**D**) Co-staining of GAD67 and β-Gal in the *Sipa1l1*^*+/-*^ hippocampal CA1 region. Arrows show *Sipa1l1* promoter activity in GABAergic neurons. (**E** and **F**) Co-staining of GAD67 and SIPA1L1 in the WT (E) or *Sipa1l1*^*-/-*^ (F) cerebral cortex. Yellow and white arrowheads indicate GAD67 positive and negative neurons, respectively. SIPA1L1 is expressed in both types of neurons. Note that dotted signals of GAD67 in the neuropil and on the surface of somata are presynaptic terminals of GABAergic neurons. so, stratum oriens; sp, stratum pyramidale; sr, stratum radiatum; slm, stratum lacunosum-moleculare. (A and B) or (C-F) are fluorescent or confocal microscopic images, respectively. Scale bars: (A) 200 μm; (B, D) 100 μm; (C, E, F) 10 μm.

**Figure S3.**
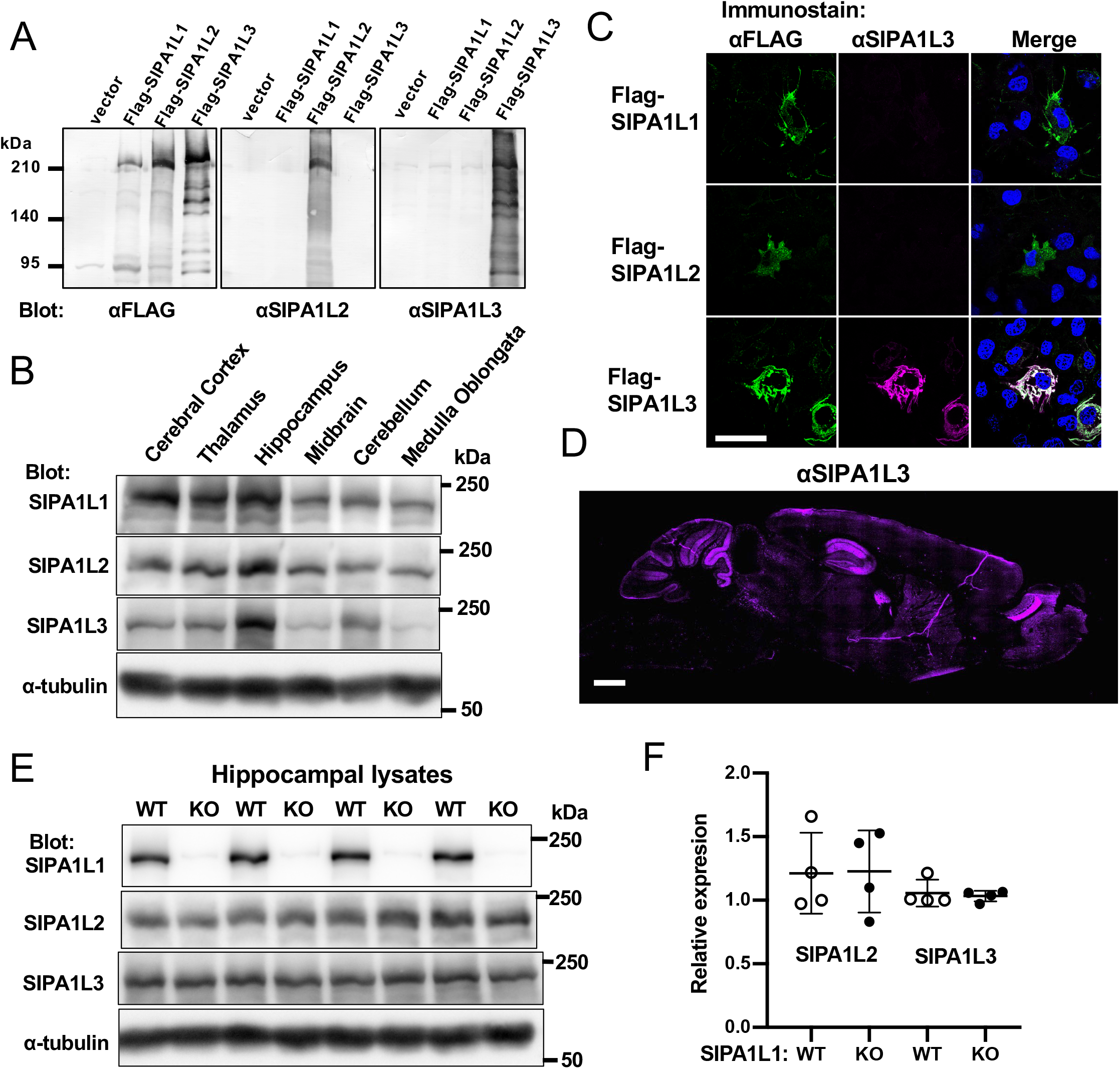
No significant up-regulation of SIPA1L1 paralogs in *Sipa1l1*^*-/-*^ hippocampus. (**A, B**) Validation of Anti-SIPA1L2 and anti-SIPA1L3 antibodies by immunoblotting (A) and immunostaining (B). The FLAG-tagged SIPA1L family proteins indicated were exogenously expressed in HEK293T (A) or COS-7 (B) cells. Anti-SIPA1L2 and anti-SIPA1L3 antibodies did not cross react with other paralogs. (**C**) Lysates of indicated regions of mature WT brain were subjected to immunoblotting. (**D**) Immunofluorescence staining of SIPA1L3 on sagittal sections from WT brain. SIPA1L2 was not detectable by immunostaining of mouse brain. (**E**) WT and KO hippocampal lysates from littermate pair were placed side by side for immunoblotting. (**F**) Quantification of (E), mean ± SD is shown. t (3) = 0.113, P = 0.92 for SIPA1L2 and t (3) = 0.419, P = 0.70 for SIPA1L3; paired two-tailed t test. Scale bars: (B) 500 μm; (D) 1000 μm

**Figure S4.**
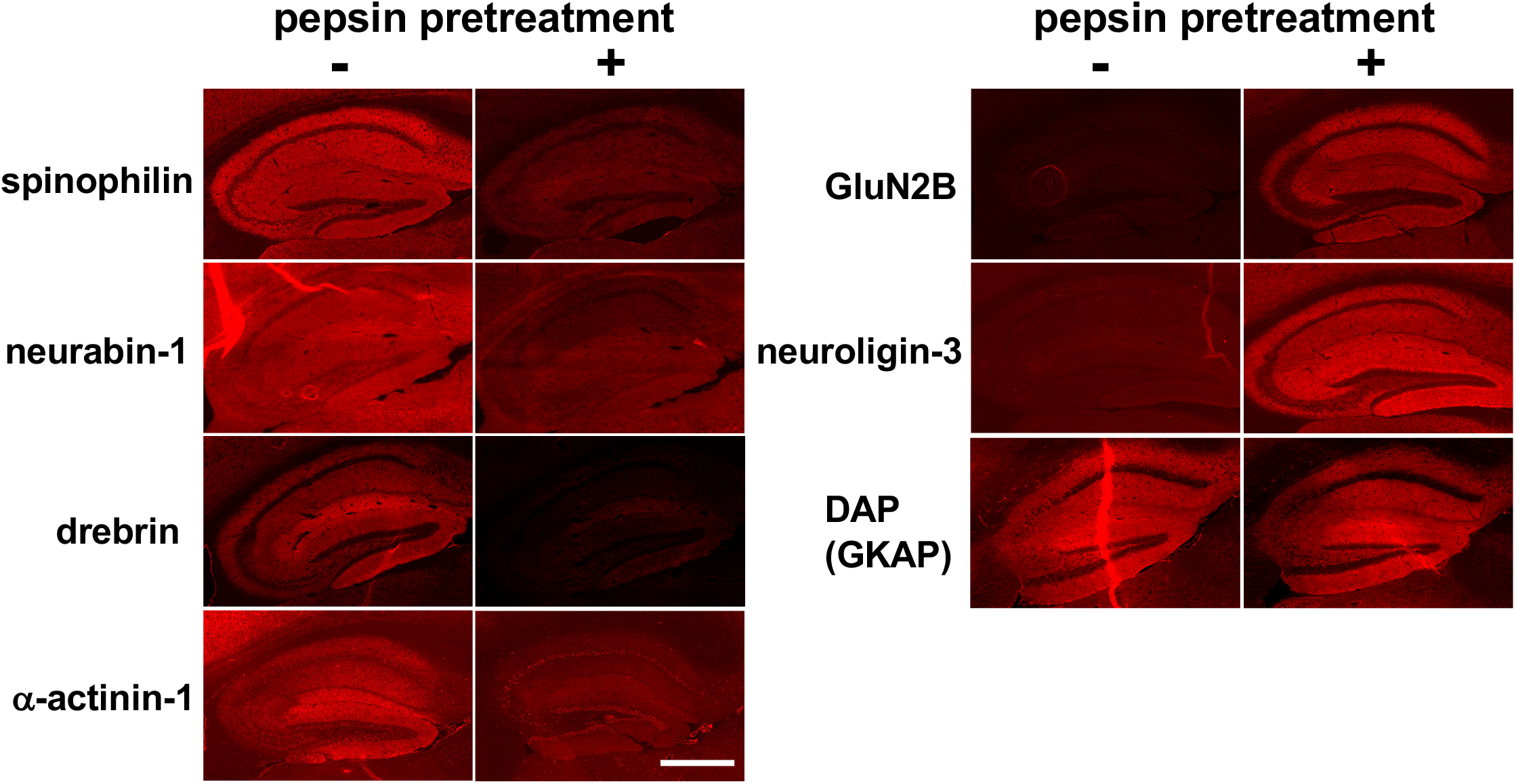
SIPA1L1-interacting proteins generally show non-PSD-like staining pattern. Immunofluorescent images of indicated proteins on pepsin-pretreated (+) or -untreated (-) hippocampal slices. Paired images were acquired under the identical settings and conditions. The PSD proteins show strong staining in pepsin-pretreated hippocampal slices (right column), whereas SIPA1L1-interacting proteins show stronger staining in untreated hippocampal slices (left column). DAP/GKAP has been shown to have both PSD and non-PSD populations (Ref. 29). Scale bar: 500 μm.

**Figure S5.**
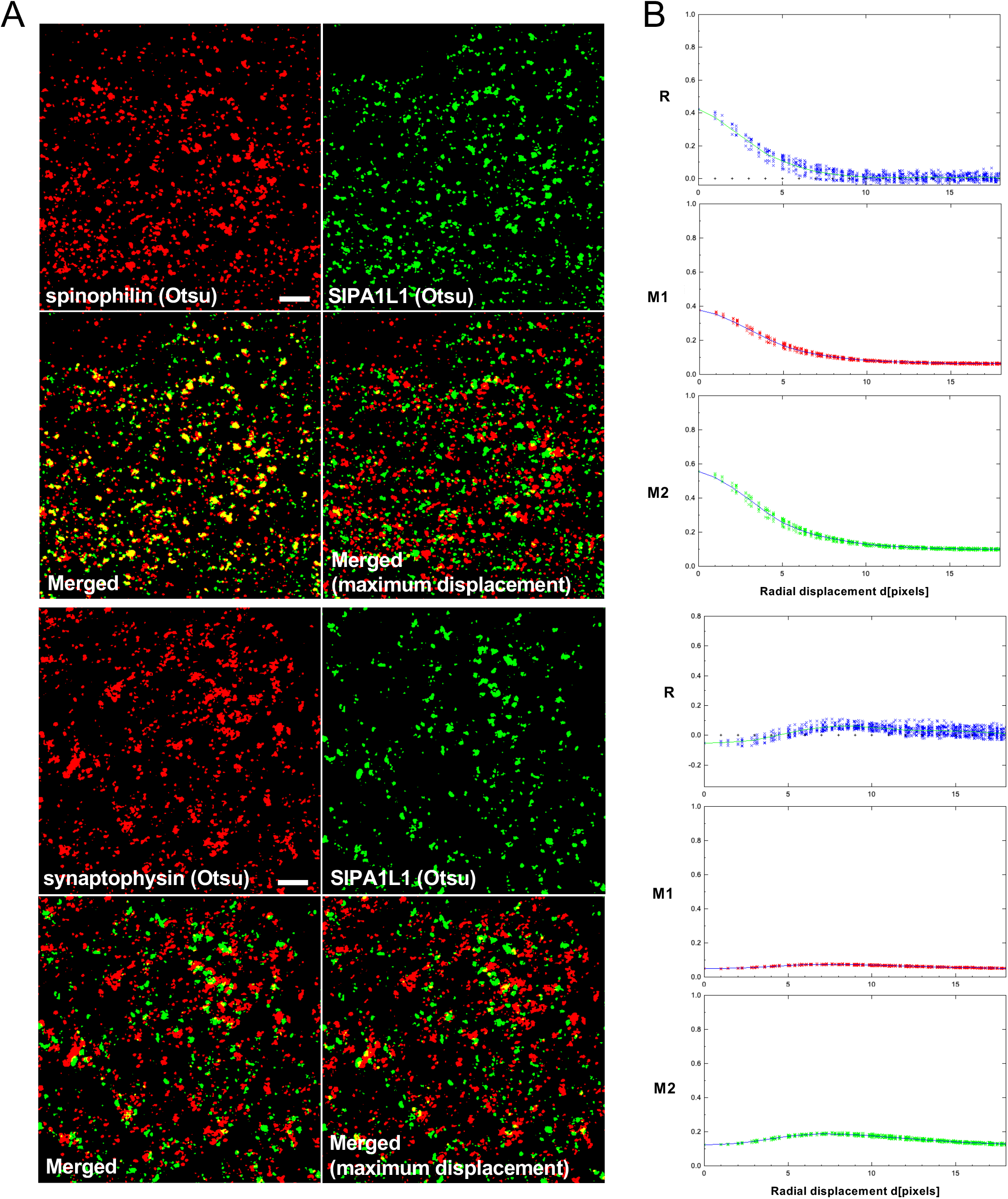
Co-localization analysis using Confined Displacement Algorithm. **(A)** Examples of super-resolution images converted by Otsu method to define background area. Otsu images were used to calculate Manders coefficient, whereas original images were used to calculate Pearson correlation coefficient. Random displacement images to calculate the statistical significance were generated by translations in all directions at set distances (see Methods for details). Scale bar: 2 μm. (**B**) R, Pearson correlation coefficient; M1, Manders coefficient for spiniophilin or synaptophysin; M2, Manders coefficient for SIPA1L1.

**Figure S6.**
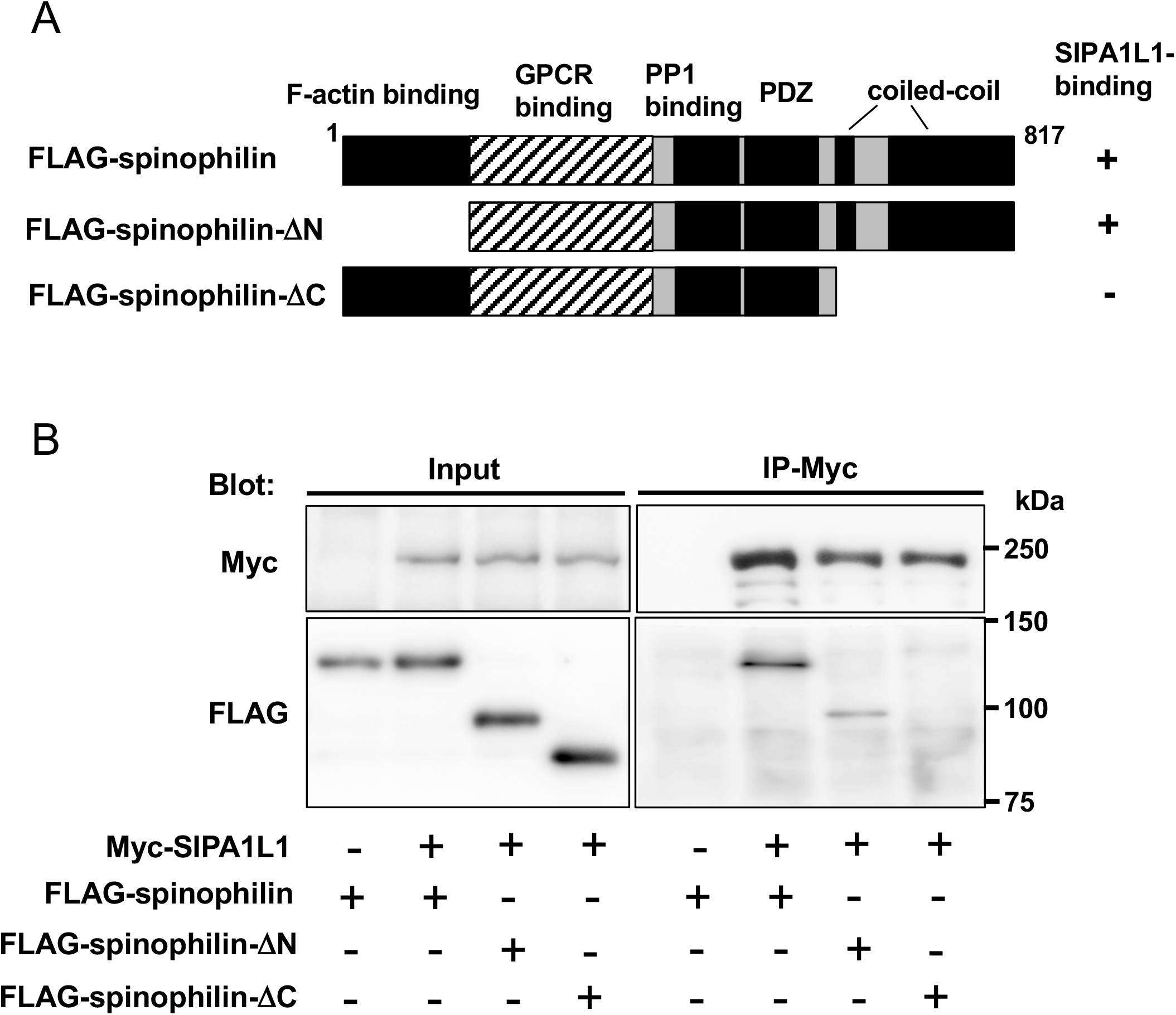
The SIPA1L1-spinophilin interaction in COS-7 cells. **(A)** Domain organization of spinophilin. A schematic structure of deletion mutants of spinophilin is shown. Positive binding activity is indicated as +. (**B**) Myc-tagged SIPA1L1 and indicated FLAG-tagged spinophilin constructs were exogenously co-expressed in COS-7 cells. The complex was immunoprecipitated with anti-Myc antibodies and interactions were examined by Western blotting. Expressed constructs are indicated by +. The interaction depended on the C-terminal coiled-coil domain, but not on the N-terminal F-actin-binding domain of spinophilin.

**Figure S7.**
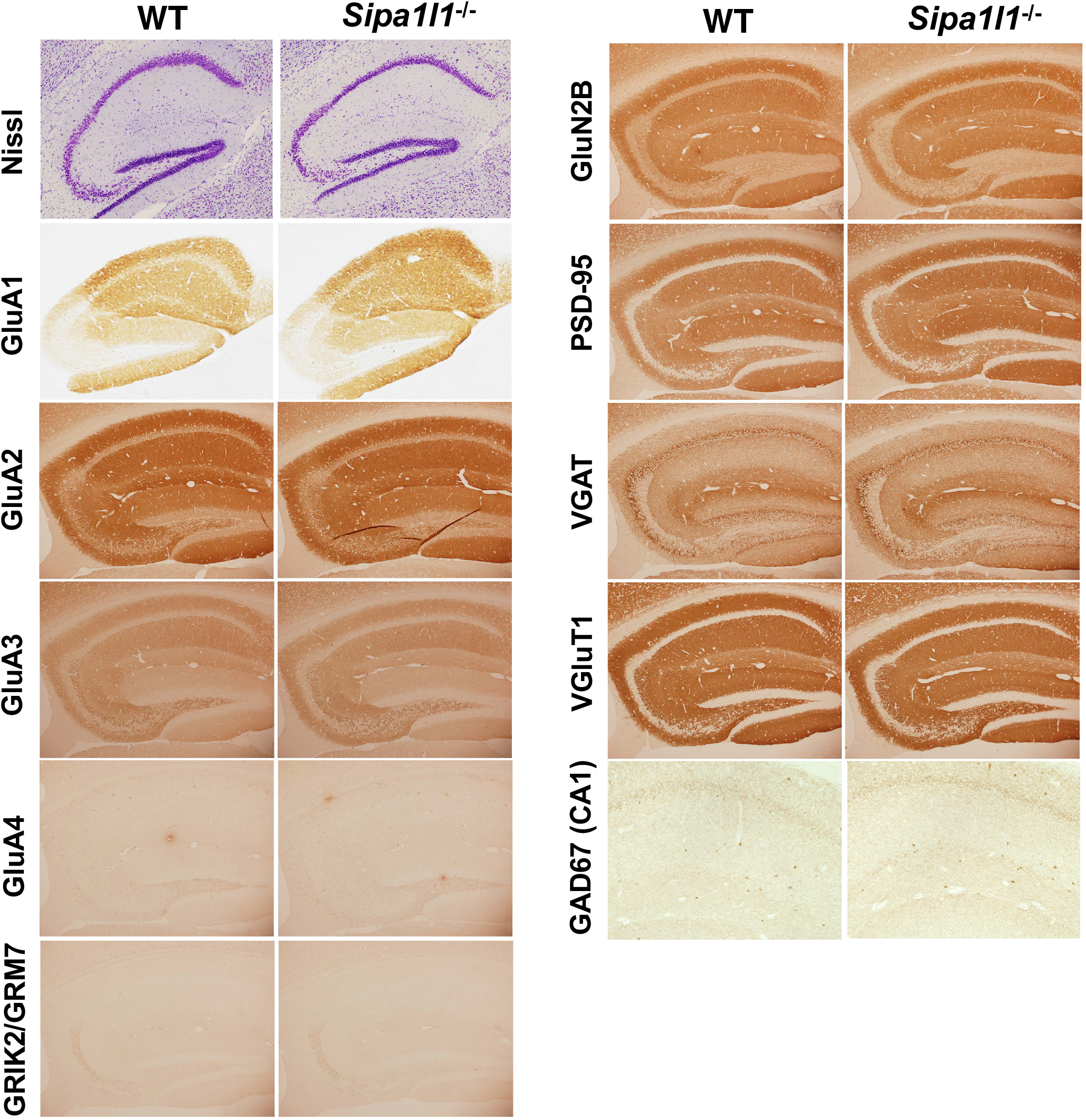
No gross defects in development of the *Sipa1l1*^*-/-*^ brain. Representative data of Nissl staining or immunostaining of synaptic marker proteins performed on adult *Sipa1l1*^*-/-*^ mouse brains and those of WT littermates.

**Table S3.**
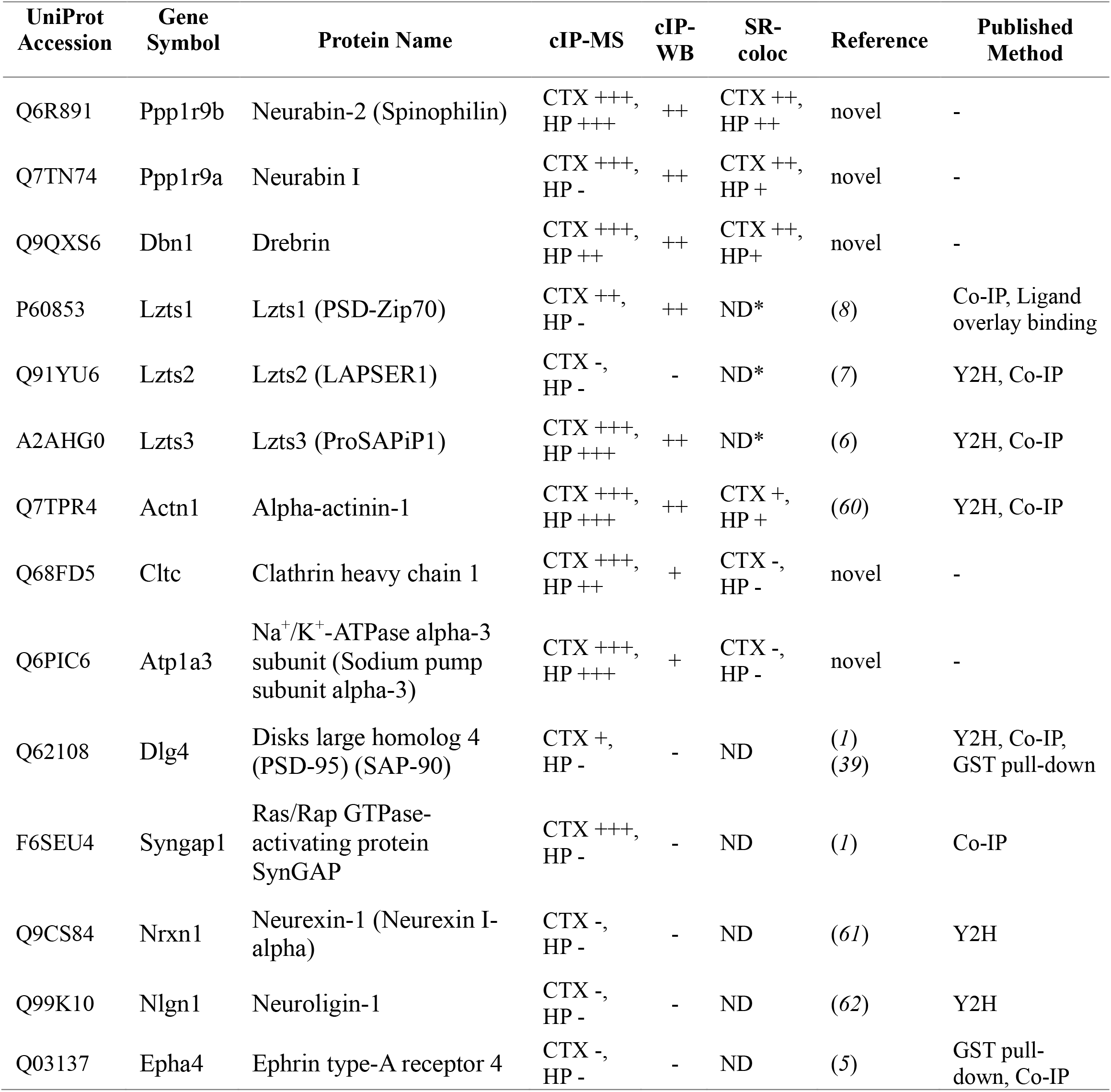
Summary of the screening results for SIPA1L1-interacting proteins. CTX, cerebral cortex; HP, hippocampus; ND, not determined; Co-IP, co-immunoprecipitation; Y2H: yeast two-hybrid screen; GST, Glutathione-S-transferase; *, not performed due to a lack of suitable antibody.

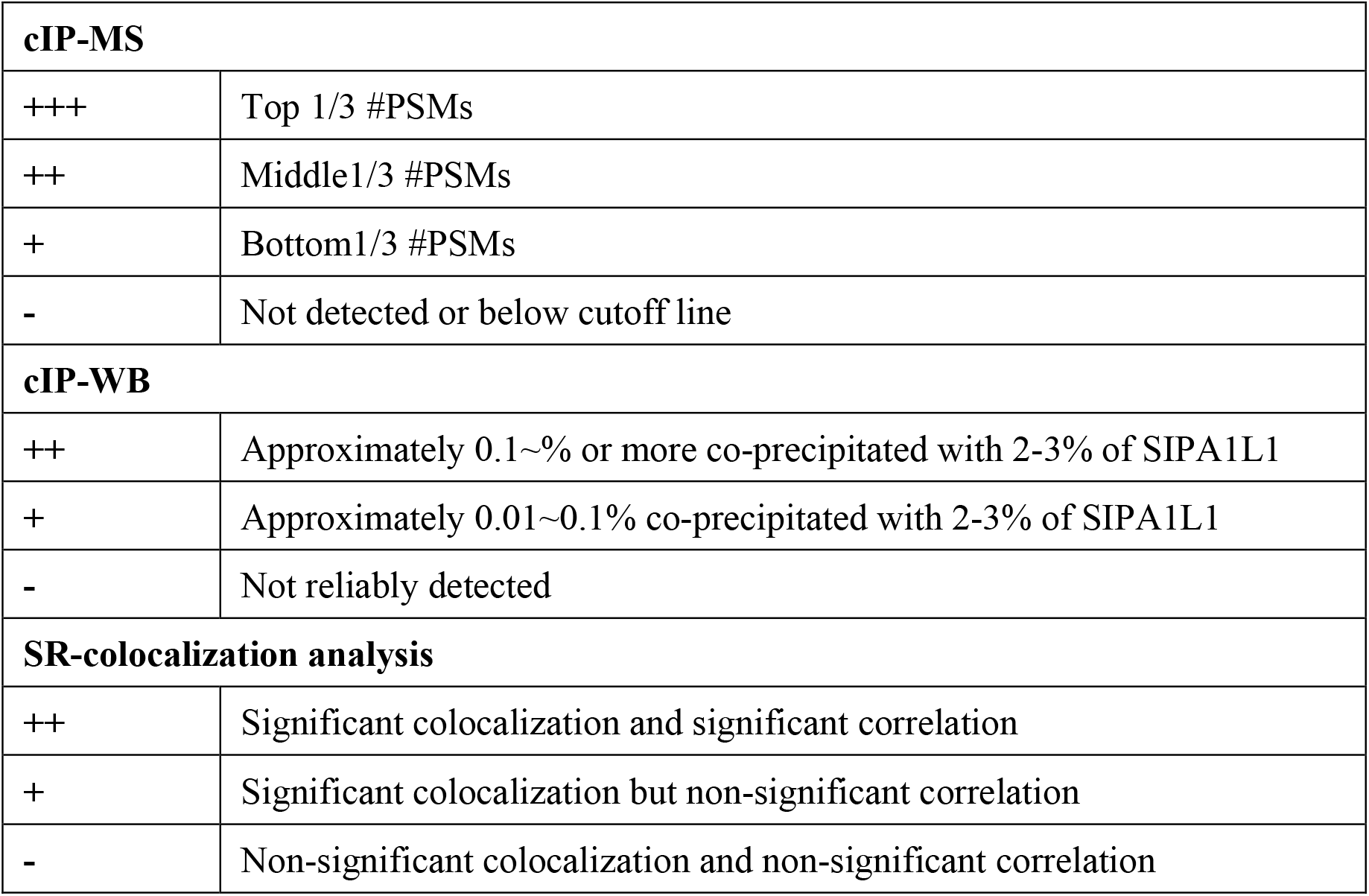

**Table S5.**
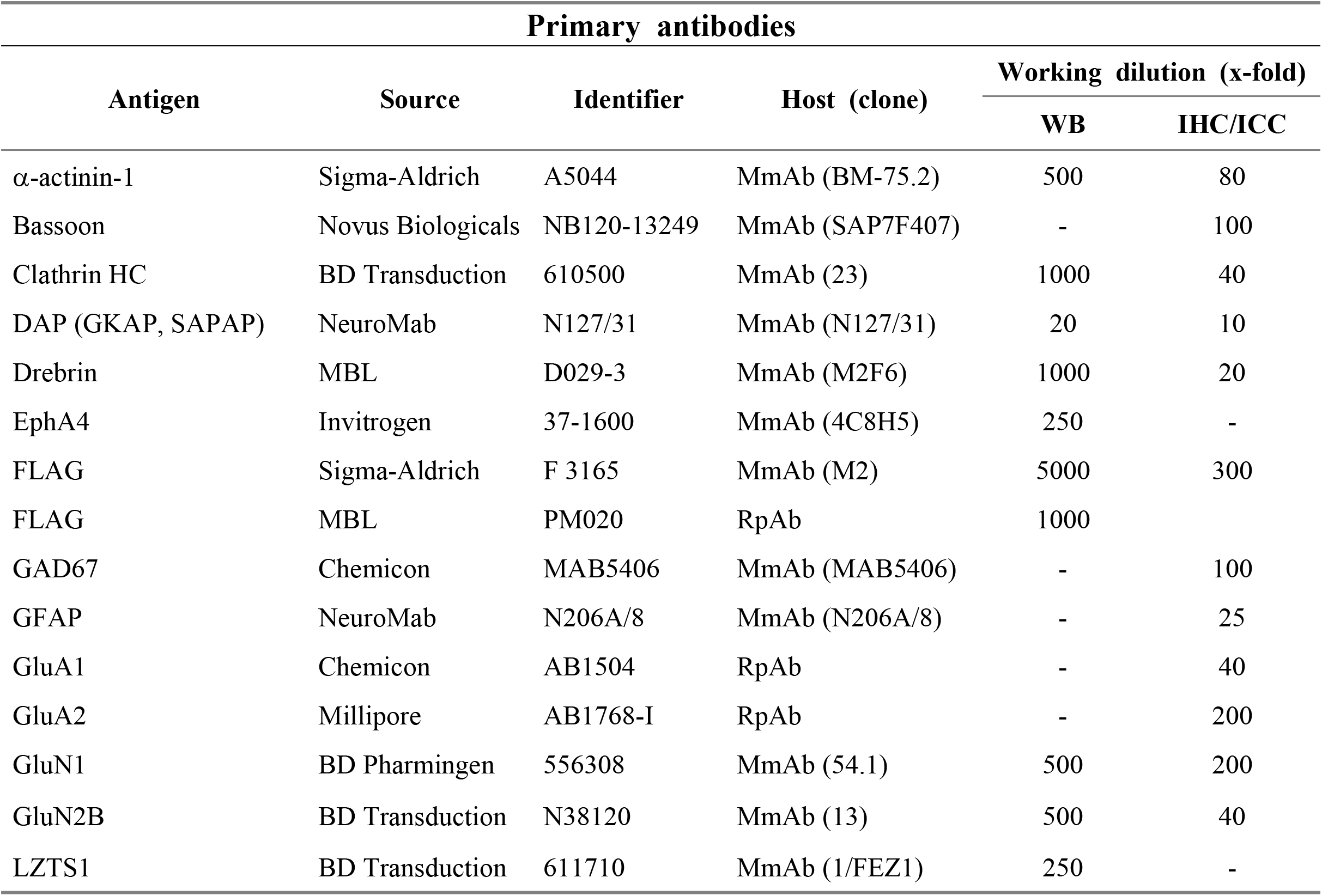
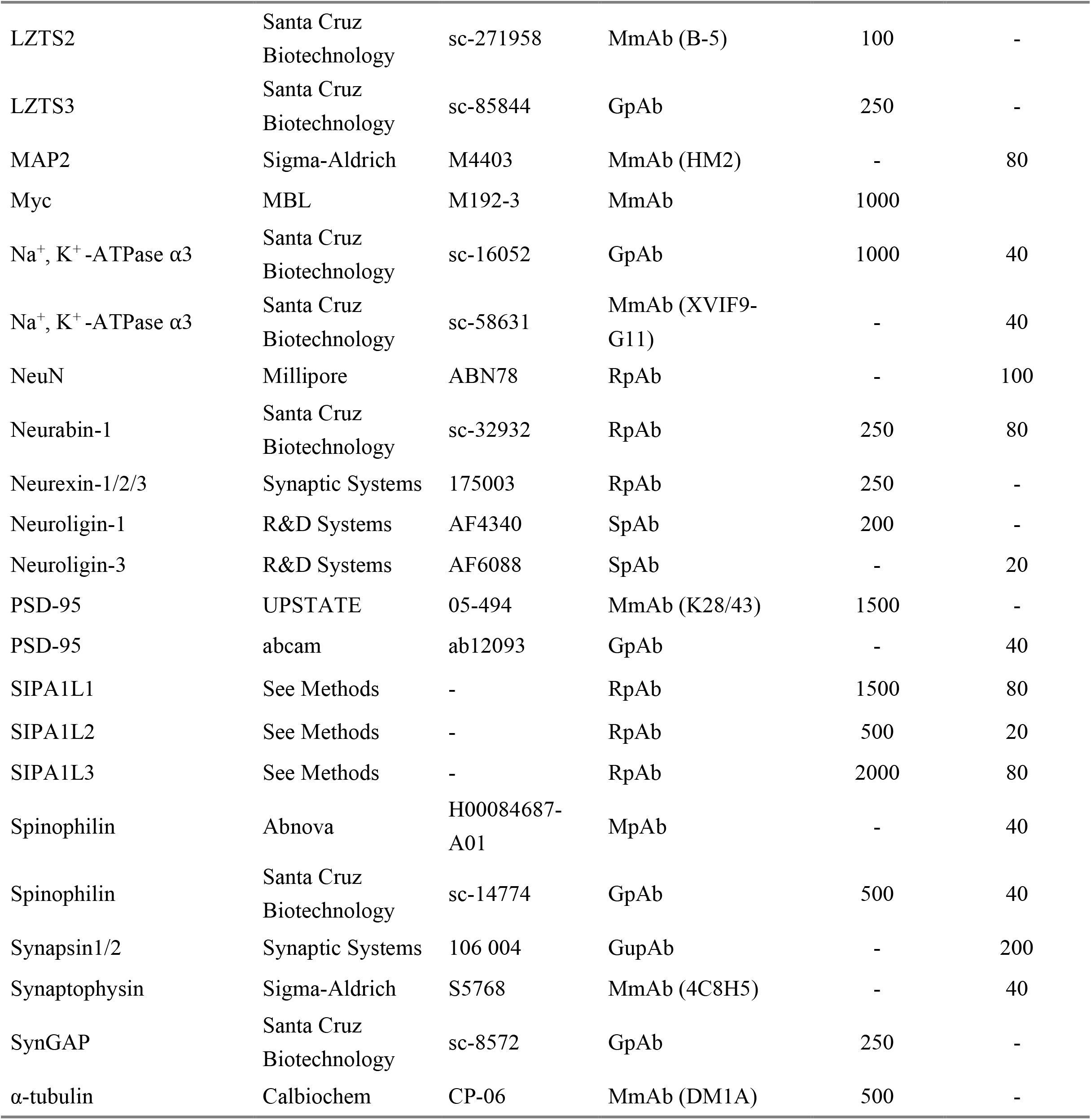
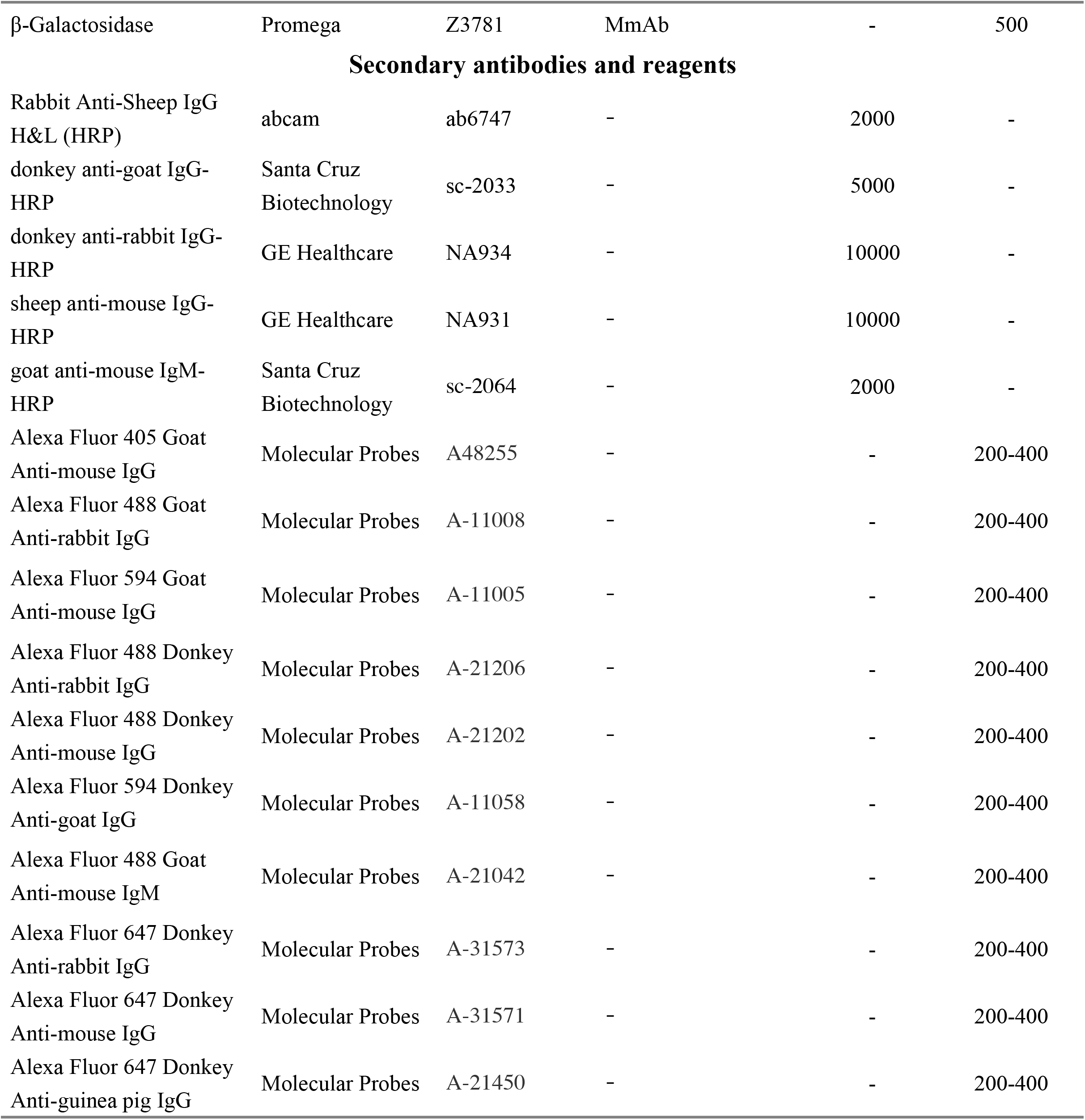
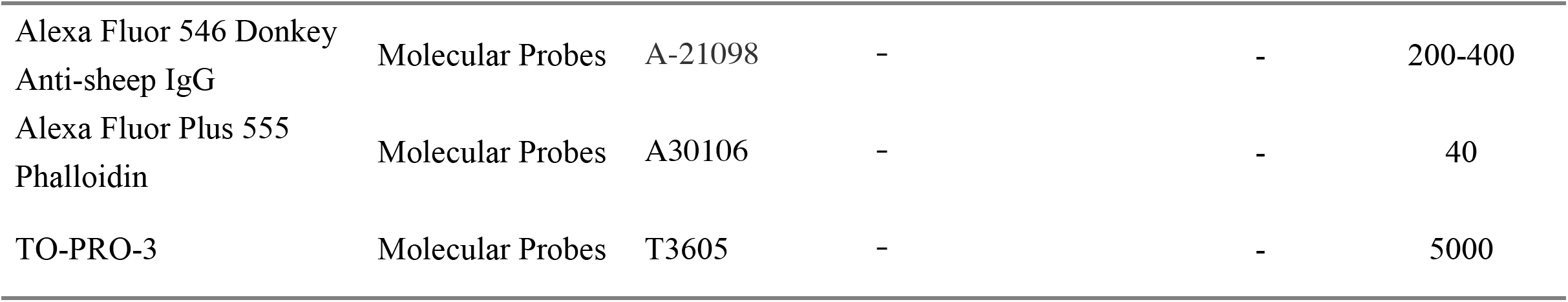
Antibodies and reagents. R, rabbit; M, mouse; G, goat; S, sheep; Gu, guinea pig; pAb, polyclonal antibody; mAb, monoclonal antibody. MpAb (Abnova) and GpAb (SantaCruz) for spinophilin, which were raised against different regions of spinophilin, both gave similar results when double stained with SIPA1L1. MmAb (SantaCruz, clone: XVIF9-G11) and GpAb (SantaCruz) for Na^+^, K^+^ -ATPase α3, which were raised against different immunogen, both gave similar results when double stained with SIPA1L1. Results from MpAb or GpAb for spinophilin or Na^+^, K^+^ -ATPase α3, respectively, are shown in the figures.

